# Macroscopic Coherent Structures in a Stochastic Neural Network: From Interface Dynamics to Coarse-Grained Bifurcation Analysis

**DOI:** 10.1101/043976

**Authors:** Daniele Avitabile, Kyle Wedgwood

## Abstract

We study coarse pattern formation in a cellular automaton modelling a spatially-extended stochastic neural network. The model, originally proposed by Gong and Robinson [36], is known to support stationary and travelling bumps of localised activity. We pose the model on a ring and study the existence and stability of these patterns in various limits using a combination of analytical and numerical techniques. In a purely deterministic version of the model, posed on a continuum, we construct bumps and travelling waves analytically using standard interface methods from neural fields theory. In a stochastic version with Heaviside firing rate, we construct approximate analytical probability mass functions associated with bumps and travelling waves. In the full stochastic model posed on a discrete lattice, where a coarse analytic description is unavailable, we compute patterns and their linear stability using equation-free methods. The lifting procedure used in the coarse time-stepper is informed by the analysis in the deterministic and stochastic limits. In all settings, we identify the synaptic profile as a mesoscopic variable, and the width of the corresponding activity set as a macroscopic variable. Stationary and travelling bumps have similar meso‐ and macroscopic profiles, but different microscopic structure, hence we propose lifting operators which use microscopic motifs to disambiguate between them. We provide numerical evidence that waves are supported by a combination of high synaptic gain and long refractory times, while meandering bumps are elicited by short refractory times.

## 1. Introduction

In the past decades, single-neuron recordings have been complemented by multineuronal experimental techniques, which have provided quantitative evidence that the cells forming the nervous systems are coupled both structurally [8] and functionally (for a recent review, see [75] and references therein). An important question in neuroscience concerns the relationship between electrical activity at the level of individual neurons and the emerging spatio-temporal coherent structures observed experimentally using local field potential recordings [22], functional magnetic resonance imaging [69] and electroencephalography [58].

There exist a wide variety of models describing activity at the level of an individual neuron [39, 26], and major research efforts in theoretical and computational neuroscience are directed towards coupling neurons in large-dimensional neural networks, whose behaviour is studied mainly via direct numerical simulations [40, 27].

A complementary approach, dating back to Wilson and Cowan [73, 74] and Amari [1, 2], foregoes describing activity at the single neuron level by representing averaged activity across populations of neurons. These *neural field models* are nonlocal, spatially-extended, excitable pattern-forming systems [24] which are often analytically tractable and support several coherent structures such as localised radially-symmetric states [72, 54, 52, 14, 29], localised patches [53, 63, 4], patterns on lattices with various symmetries [23, 13], travelling bumps and fronts [25, 12], rings [61, 19], breathers [30, 31, 32], target patterns [20], spiral waves [47] and lurching waves [35, 60, 71] (for comprehensive reviews, we refer the reader to [11, 12]).

Recent studies have analysed neural fields with additive noise [38, 28, 46], multiplicative noise [15], or noisy firing thresholds [7], albeit these models are still mostly phenomenological. Even though several papers derive continuum neural fields from microscopic models of coupled neurons [41, 9, 10, 5], the development of a rigorous theory of multi-scale brain models is an active area of research.

Numerical studies of networks based on realistic neural biophysical models rely almost entirely on brute-force Monte Carlo simulations (for a very recent, remarkable example, we refer the reader to [56]). With this *direct numerical simulation* approach, the stochastic evolution of each neuron in the network is monitored, resulting in huge computational costs, both in terms of computing time and memory. From this point of view, multi-scale numerical techniques for neural networks present interesting open problems.

When few clusters of neurons with similar properties form in the network, a significant reduction in computational costs can be obtained by population density methods [59, 37], which evolve probability density functions of neural subpopulations, as opposed to single neuron trajectories. This coarse-graining technique is particularly effective when the underlying microscopic neuronal model has a low-dimensional state space (such as the leaky integrate-and-fire model) but its performance degrades for more realistic biophysical models. Developments of the population density method involve analytically derived moment closure approximations [16, 55]. Both Monte Carlo simulations and population density methods give access only to stable asymptotic states, which may form only after long-transient simulations.

An alternative approach is offered by *equation-free* [42, 43] and *heterogeneous multiscale* methods [70, 21], which implement multiple-scale simulations using an on-the-fly numerical closure approximations. Equation-free methods, in particular, are of interest in computational neuroscience as they accelerate macroscopic simulations and allow the computation of unstable macroscopic states. In addition, with equation-free methods, it is possible to perform coarse-grained bifurcation analysis using standard numerical bifurcation techniques for time-steppers [68].

The equation-free framework [42, 43] assumes the existence of a closed coarse model in terms of a few macroscopic state variables. The model closure is enforced numerically, rather than analytically, using a *coarse time-stepper*: a computational procedure which takes advantage of knowledge of the microscopic dynamics to time-step an approximated macroscopic evolution equation. A single coarse time step from time *t*_0_ to time *t*_1_ is composed of three stages: (i) *lifting*, that is, the creation of microscopic initial conditions that are compatible with the macroscopic states at time *t*_0_; (ii) *evolution*, the use of independent realisations of the microscopic model over a time interval [*t*_0_, *t*_1_]; (iii) *restriction*, that is, the estimation of the macroscopic state at time *t*_1_ using the realisations of the microscopic model.

While equation-free methods have been employed in various contexts (see [43] and references therein) and in particular in neuroscience applications [48, 50, 49, 66, 67, 51], there are still open questions, mainly related to how noise propagates through the coarse time stepper. A key aspect of every equation-free implementation is the lifting step. The underlying lifting operator, which maps a macroscopic state to a set of microscopic states, is generally non-unique, and lifting choices have a considerable impact on the convergence properties of the resulting numerical scheme [3]. Even though the choice of coarse variables can be automatised using data-mining techniques, as shown in several papers by Laing, Kevrekidis and co-workers [48, 50, 49], the lifting step is inherently problem dependent.

The present paper explores the possibility of using techniques from neural field models to inform the coarse-grained bifurcation analysis of discrete neural networks. A successful strategy in analysing neural fields is to replace the models' sigmoidal firing rate functions with Heaviside distributions [11, 12]. Using this strategy, it is possible to relate macroscopic observables, such as bump widths or wave speeds, to biophysical parameters, such as firing rate thresholds. Under this hypothesis, a macroscopic variable suggests itself, as the state of the system can be constructed entirely via the loci of points in physical space where the neural activity attains the firing-rate threshold value. In addition, there exists a closed (albeit implicit) evolution equation for such interfaces [19].

In this study, we show how the insight gained in the Heaviside limit may be used to perform coarse-grained bifurcation analysis of neural networks, even in cases where the network does not evolve according to an integro-differential equation. As an illustrative example, we consider a spatially-extended neural network in the form of a discrete time Markov chain with discrete ternary state space, posed on a lattice. The model is an existing cellular automaton proposed by Gong and co-workers [36], and it has been related to neuroscience in the context of relevant spatio-temporal activity patterns that are observed in cortical tissue. In spite of its simplicity, the model possesses sufficient complexity to support rich dynamical behaviour akin to that produced by neural fields. In particular, it explicitly includes refractoriness and is one of the simplest models capable of generating propagating activity in the form of travelling waves. An important feature of this model is that the microscopic transition probabilities depend on the local properties of the tissue, as well as on the global synaptic profile across the network. The latter has a convolution structure typical of neural field models, which we exploit to use interface dynamics and define a suitable lifting strategy.

We initially study the model in simplifying limits in which an analytical (or semi-analytical) treatment is possible. In these cases, we construct bump and wave solutions and compute their stability. This analysis follows the standard Amari framework, but is here applied directly to the cellular automaton. We then derive the corresponding lifting operators, which highlight a critical importance of the microscopic structure of solutions: one of the main results of our analysis is that, since macroscopic stationary and travelling bumps coexist and have nearly identical macroscopic profiles, a standard lifting is unable to distinguish between them, thereby preventing coarse numerical continuation. These structures, however, possess different microstructures, which are captured by our analysis and subsequently by our lifting operators. This allows us to compute separate solution branches, in which we vary several model parameters, including those associated with the noise processes.

The manuscript is arranged as follows: In Section 2 we outline the model. In Section 3, we simulate the model and identify the macroscopic profiles in which we are interested, together with the coarse variables that describe them. In Section 4, we define a deterministic version of the full model and lay down the framework for analysing it. In Sections 5 and 6, we respectively construct bump and wave solutions under the deterministic approximation and compute the stability of these solutions. In Section 7, we define and construct travelling waves relaxing the deterministic limit. In Sections 8.1 and 8.2, we provide the lifting steps for use in the equation-free algorithm for the bump and wave respectively. In Section 9, we briefly outline the continuation algorithm and in Section 10, we show the results of applying this continuation to our system. Finally, in Section 11, we make some concluding remarks.

## 2. Model description

### 2.1. State variables for continuum and discrete tissues

In this section, we present a modification of a model originally proposed by Gong and Robinson [36].

We consider a one-dimensional neural tissue 𝕏 ⊂ ℝ. At each discrete time step *t* ∈ ℤ, a neuron at position *x* ∈ 𝕏 may be in one of three states: a refractory state (henceforth denoted as –1), a quiescent state (0) or a spiking state (1). Our state variable is thus a function *u* : 𝕏 × → 𝕌, where 𝕌 = {–1, 0, 1}. We pose the model on a continuum tissue 𝕊 = ℝ/2*L*ℤ or on a discrete tissue featuring *N* + 1 evenly spaced neurons,

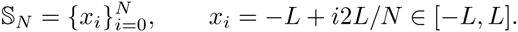

We will often alternate between the discrete and the continuum setting, hence we will use a unified notation for these cases. We use the symbol *X* to refer to either 𝕊 or 𝕊_*N*_, depending on the context. Also, we use *u*(·, *t*) to indicate the state variable in both the discrete and the continuum case: *u*(·, *t*) will denote a step function defined on 𝕊 in the continuum case and a vector in 𝕌 ^*N*^ with components *u*(*x*_*i*_, *t*) in the discrete case. Similarly, we write ∫_𝕏_ *u*(*x*) d*x* to indicate ∫_𝕊_ *u*(*x*) d*x* or 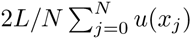.

### 2.2. Model definition

We use the term *stochastic model* when the Markov chain model described below is posed on 𝕊_*N*_. An example of microscopic state supported by the stochastic model is given in Figure 1(a).

**Fig. 1.**
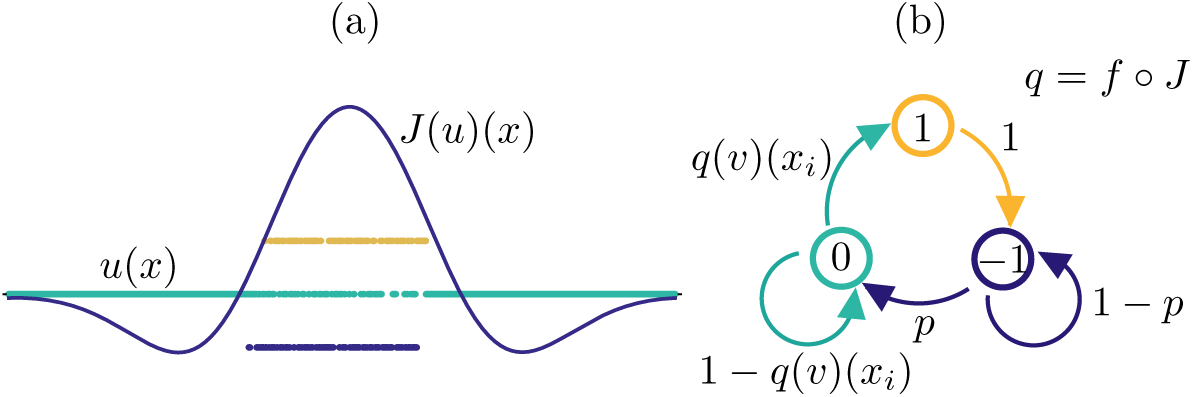
*(a)*: Example of microscopic state *u*(*x*) ∈ 𝕌 ^*N*^ and corresponding synaptic profile *J*(*u*)(*x*) ∈ ℝ^*N*^ in a network of 1024 neurons, (b): Schematic of the transition kernel for the network (see also Equations (2.5)–(2.7)), The conditional probability of the local variable *u*(*x*_*i*_, *t* + 1) depends on the global state of the network at time *t*, via the function *q* = *f* ∘ *J*, as seen in (2.7).

In the model, neurons are coupled via a translation-invariant synaptic kernel, that is, we assume the connection strength between two neurons to be dependent on their mutual Euclidean distance. In particular, we prescribe that short range connections are excitatory, whilst long-range connections are inhibitory. To model this coupling, we use a standard Mexican hat function,

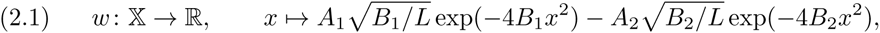

and denote by *W* its periodic extension.

In order to describe the dynamics of the model, it is useful to partition the tissue 𝕏 into the 3 pullback sets

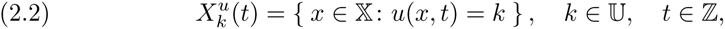

so that we can write, for instance, 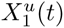 to denote the set of neurons that are firing at time *t* (and similarly for 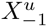 and 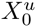). Where it is unambiguous, we shall simply write *X*_*k*_ or *X*_*k*_(*t*) in place of 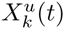.

The synaptic input to a cell at position *x*_*i*_ is given by a weighted sum of inputs from all firing cells. Using the synaptic kernel (2.1) and the partition (2.2), the synaptic input is then modelled as

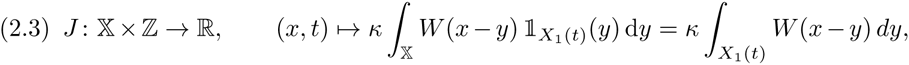

where *κ* ∈ ℝ_+_ is the synaptic gain, which is common for all neurons and 𝟙_*X*_ is the indicator function of a set *X*.

#### Remark 2.1 (Synaptic input as mesoscopic variable).

Since *X*_1_ depends on the microscopic state variable u, so does the synaptic input (2.3). Where necessary, we will write *J*(*u*)(*x*, *t*) to highlight the dependence on *u*. We refer the reader to Figure 1 for a concrete example of synaptic profile. It will become apparent in the next section that *J* plays the role of a mesoscopic state variable associated with u.

The firing probability associated to a quiescent neuron is linked to the synaptic input via the firing rate function

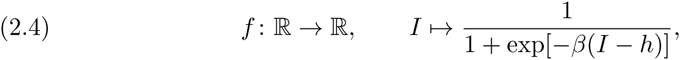

whose steepness and threshold are denoted by the positive real numbers *β* and *h*, respectively. We are now ready to describe the evolution of the stochastic model, which is a discrete-time Markov process with finite state space 𝕌 ^*N*^ and transition probabilities specified as follows: for each *x*_*i*_ ∈ 𝕊_*N*_ and *t* ∈ ℤ

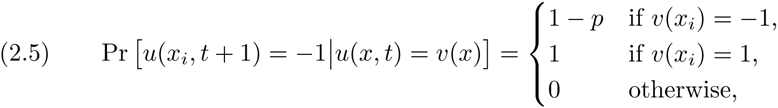

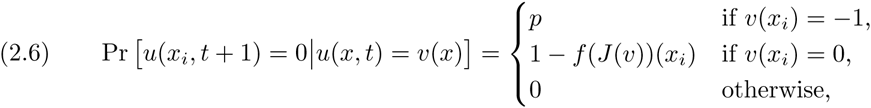

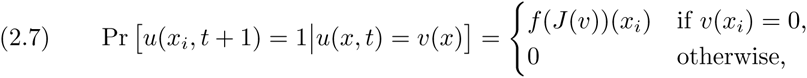

where *p* ∈ (0,1]. We give a schematic representation of the transitions of each neuron in the network in Figure 1(b). We remark that conditional probability of the *local* variable *u*(*x*_*i*_, *t* + 1) depends on the *global* state of the network at time *t*, via the function *f* ∘ *J*.

The model described by (2.1)–(2.7), complemented by initial conditions, defines a stochastic evolution map that we will formally denote as

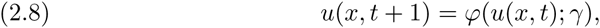

where *γ* = (*κ*, *β*, *h*, *p*, *A*_1_, *A*_2_, *B*_1_, *B*_2_) is a vector of control parameters.

## 3. Microscopic states observed via direct simulation

In this section, we introduce a few coherent states supported by the stochastic model. The main aim of the section is to show examples of bumps, multiple bumps and travelling waves, whose existence and stability properties will be studied in the following sections. In addition, we give a first characterisation of the macroscopic variables of interest and link them to the microscopic structure observed numerically.

### 3.1. Bumps

In a suitable region of parameter space, the microscopic model supports bump solutions [62] in which the microscopic variable *u*(*x*, *t*) is nonzero only in a localised region of the tissue. In this *active* region, neurons attain all values in 𝕌. In Figure 2, we show a time simulation of the microscopic model with *N* = 1024 neurons. At each time *t*, neurons are in the refractory (blue), quiescent (green) or spiking (yellow) state. We prescribe the initial condition by setting *u*(*x*_*i*_, 0) = 0 outside of a localised region, in which *u*(*x*_*i*_, 0) are sampled randomly from 𝕌. After a short transient, a stochastic *microscopic* bump is formed. As expected due to the stochastic nature of the system [45], the active region wanders while remaining localised. A space-time section of the active region reveals a characteristic random microstructure (see Figure 2(a)). By plotting *J*(*x*, *t*), we see that the active region is well approximated by the portion of the tissue *X*_≥_ = [*ξ*_1_, *ξ*_2_] where *J* lies above the threshold *h*. A quantitative comparison between *J*(*x*, 50) and *u*(*x*, 50) is made in Figure 2(a). We interpret *J* as a *mesoscopic* variable associated with the bump, and *ξ*_1_ and *ξ*_2_ as corresponding *macroscopic* variables.

**Fig. 2.**
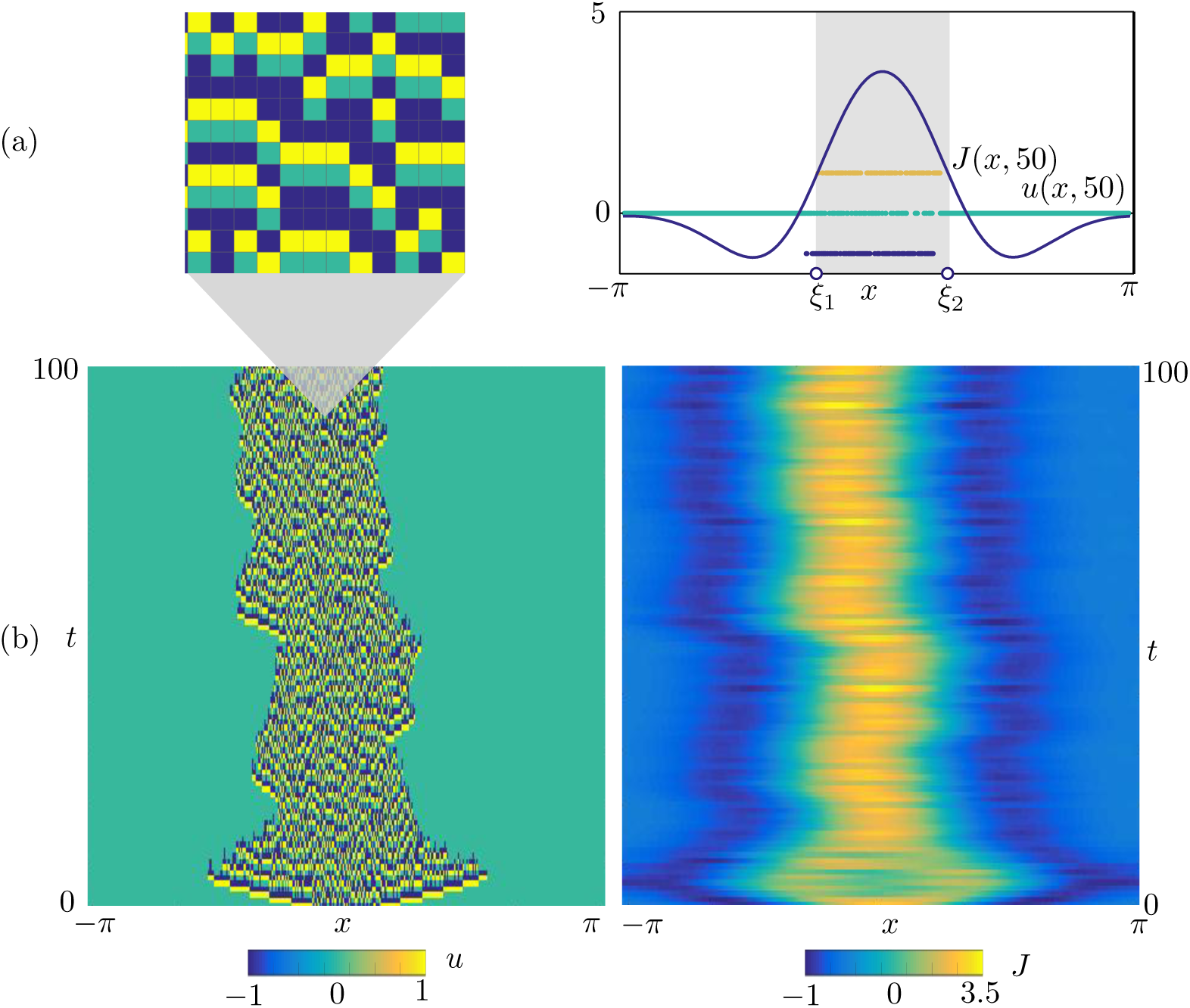
Bump obtained via time simulation of the stochastic model for (*x*, *t*) ∈ [–*π*, *π*] × [0, 100]. (a): The microscopic state *u*(*x*, *t*) (left) attains the discrete values –1 (blue), 0 (green) and 1 (yellow). The corresponding synaptic profile *J*(*x*, *t*) is a continuous function. A comparison between *J*(*x*, 50) and *u*(*x*, 50) is reported on the right panel, where we also mark the interval [*ξ*_1_, *ξ*_2_] where *J* is above the firing threshold *h*. (b): Space-time plots of *u* and *J*. Parameters as in Table 1.

### 3.2. Multiple-bumps solutions

Solutions with multiple bumps are also observed by direct simulation, as shown in Figure 3. The microstructure of these patterns resembles the one found in the single bump case (see Figure 3(a)). At the mesoscopic level, the set for which *J* lies above the threshold h is now a union of disjoint intervals [*ξ*_1_, *ξ*_2_], &, [*ξ*_7_, *ξ*_8_]. The number of bumps of the pattern depends on the width of the tissue; the experiment of Figure 3 is carried out on a domain twice as large as that of Figure 2. The examples of bump and multiple-bump solutions reported in these figures are obtained for different values of the main control parameter *κ* (see Table 1), however, these states coexist in a suitable region of parameter space, as will be shown below.

**Fig. 3.**
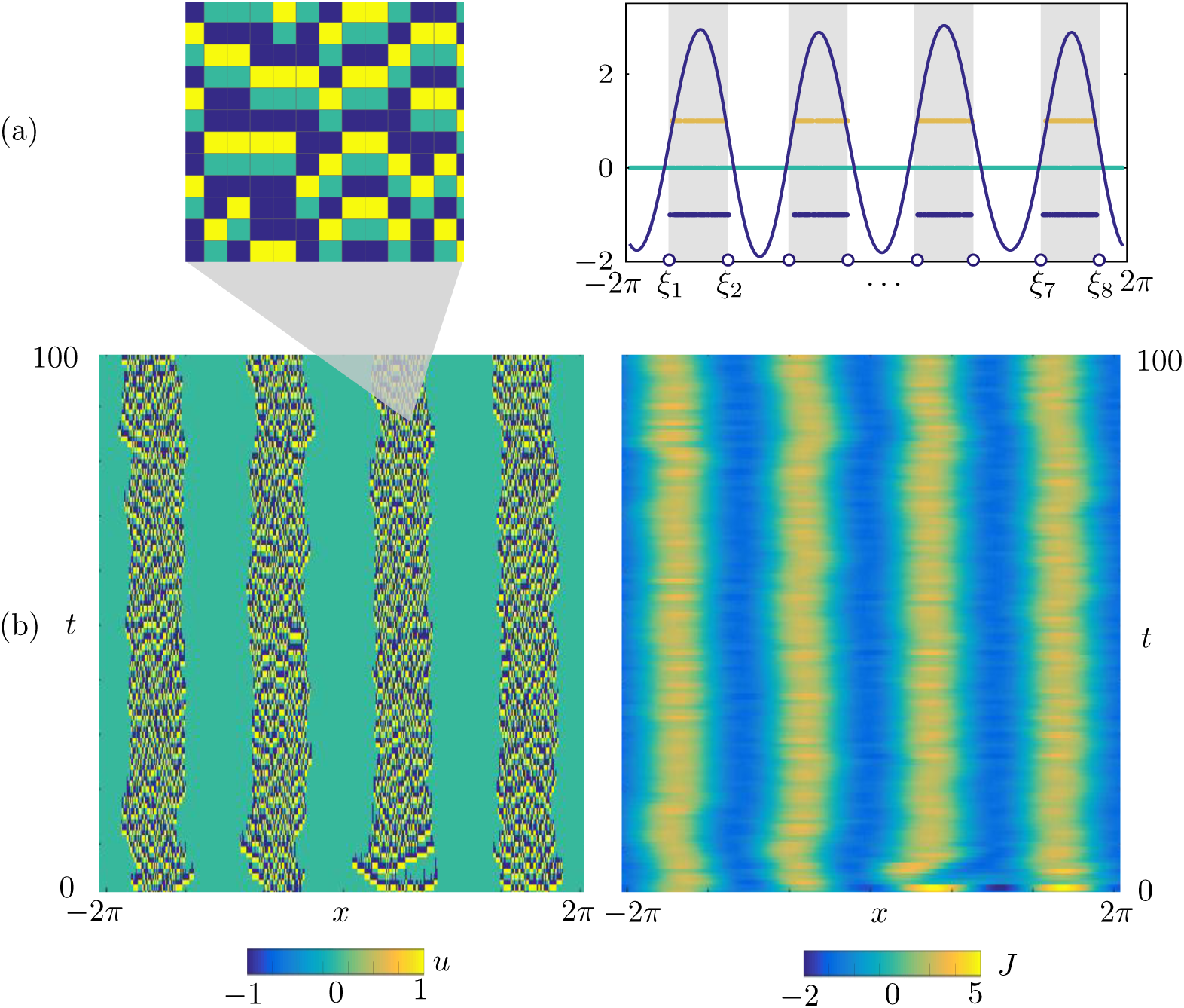
Multiple bump solution obtained via time simulation of the stochastic model for (*x*, *t*) ∈ [–2*π*, 2*π*] × [0,100], (a): The microscopic state *u*(*x*, *t*) in the active region (left) is similar to the one found for the single bump (see Figure 2(a)), A comparison between *J*(*x*, 50) and *u*(*x*, 50) is reported on the right panel, where we also mark the intervals [*ξ*_1_, *ξ*_2_],…, [*ξ*_7_, *ξ*_8_] where *J* is above the firing threshold *h*, (b): Space-time plots of *u* and *J*, Parameters are as in Table 1.

**Table 1.**
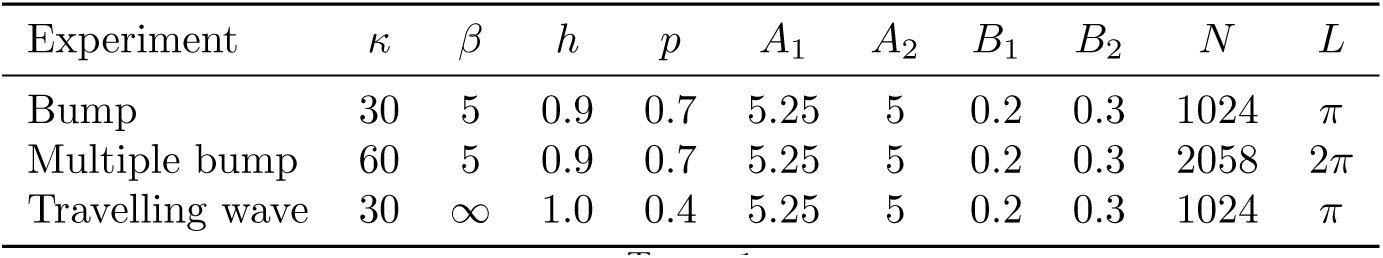
Parameter values for which the stochastic model supports a bump (Figure 2), a multiple-bump solution (Figure 3) and a travelling wave (Figure 4), The value *∞* for the parameter *β* indicates that a Heaviside firing rate has been used in place of the sigmoidal function (2.4).

| Experiment | $\kappa$ | $\beta$ | $h$ | $p$ | $A_1$ | $A_2$ | $B_1$ | $B_2$ | $N$ | $L$ |
| --- | --- | --- | --- | --- | --- | --- | --- | --- | --- | --- |
| Bump | 30 | 5 | 0.9 | 0.7 | 5.25 | 5 | 0.2 | 0.3 | 1024 | $\pi$ |
| Multiple bump | 60 | 5 | 0.9 | 0.7 | 5.25 | 5 | 0.2 | 0.3 | 2058 | $2\pi$ |
| Travelling wave | 30 | ∞ | 1.0 | 0.4 | 5.25 | 5 | 0.2 | 0.3 | 1024 | $\pi$ |

### 3.3. Travelling waves

Further simulation shows that the model also supports coherent states in the form of stochastic travelling waves. In two spatial dimensions, the system is known to support travelling spots [36, 62]. In Figure 4, we show a time simulation of the stochastic model with initial condition

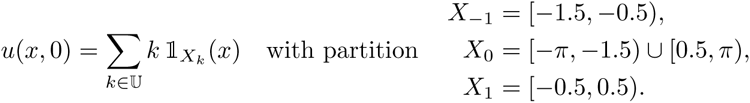

In passing, we note that the state of the network at each discrete time *t* is defined entirely by the partition {*X*_*k*_} of the tissue; we shall often use this characterisation in the reminder of the paper.

**Fig. 4.**
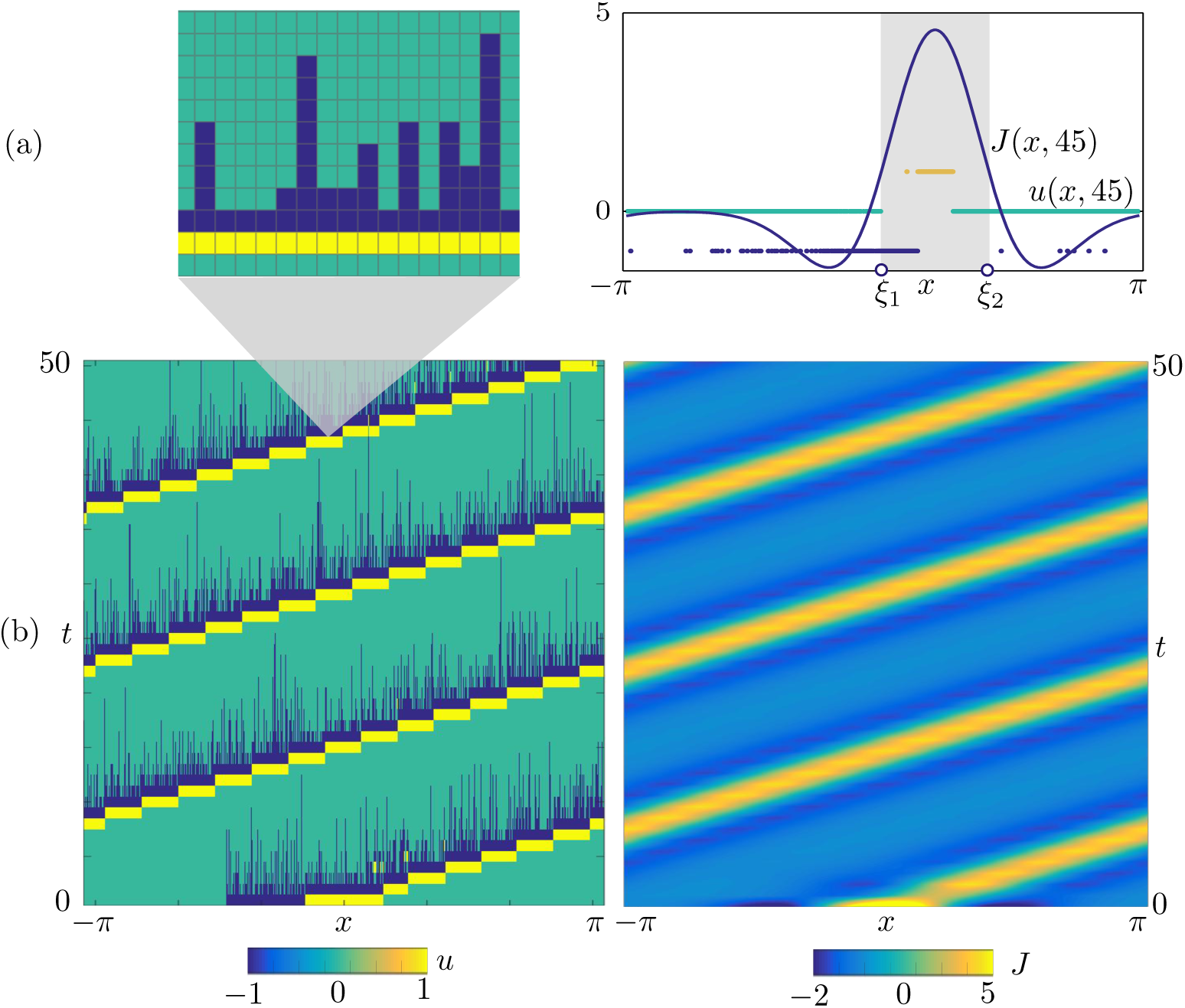
Travelling wave obtained via time simulation of the stochastic model for (*x*, *t*) ∈ [–*π*, *π*] × [0, 50]. (a): The microscopic state *u*(*x*, *t*) (left) has a characteristic microstructure, which is also visible on the right panel, where we compare *J*(*x*, 45) and *u*(*x*, 45). As in the other cases, we mark the interval [*ξ*_1_, *ξ*_2_] where *J* is above the firing threshold *h*. (b): Space-time plots of *u* and *J*. Parameters are as in Table 1.

In the direct simulation of Figure 4, the active region moves to the right and, after just 4 iterations, a travelling wave emerges. The microscopic variable, *u*(*x*, *t*), displays stochastic fluctuations which disappear at the level of the mesoscopic variable, *J*(*x*, *t*), giving rise to a seemingly deterministic travelling wave. A closer inspection (Figure 4(a)) reveals that the state can still be described in terms of the active region [*ξ*_1_, *ξ*_2_] where *J* is above *h*. However, the travelling wave has a different microstructure with respect to the bump. Proceeding from right to left, we observe:

1. A region of the tissue ahead of the wave, *x* ∈ (*ξ*_2_, *π*), where the neurons are in the quiescent state 0 with high probability.
2. An active region *x* ∈ [*ξ*_1_, *ξ*_2_], split in three subintervals, each of approximate width (*ξ*_2_ – *ξ*_1_)/3, where *u* attains with high probability the values 0, 1 and –1 respectively.
3. A region at the back of the wave, *x* ∈ [–*π*, *ξ*_1_), where neurons are either quiescent or refractory. We note that *u* = 0 with high probability as *x* → –*π* whereas, as *X* → *ξ*_1_, neurons are increasingly likely to be refractory, with *u* = –1.

A further observation of the space-time plot of *u* in Figure 4(b) reveals a remarkably simple advection mechanism of the travelling wave, which can be fully understood in terms of the transition kernel of Figure 1(b) upon noticing that, for sufficiently large *β*, *q*_*i*_ = *f*(*J*(*u*))(*x*_*i*_) ≈ 0 everywhere except in the active region, where *q*_*i*_ ≈ 1. In Figure 5, we show how the transition kernel simplifies inside and outside the active region and provide a schematic of the advection mechanism. For an idealised travelling wave profile at time *t*, we depict 3 subintervals partitioning the active region (shaded), together with 2 adjacent intervals outside the active region. Each interval is then mapped to another interval, following the simplified transition rules sketched in Figure 5(a):

1. At the front of the wave, to the right of *ξ*_2_(*t*), neurons in the quiescent state 0 remain at 0 (rules for *x* ∉ [*ξ*_1_, *ξ*_2_]).
2. Inside the active region, to the left of *ξ*_2_(*t*), we follow the rules for *x* ∈ [*ξ*_1_, *ξ*_2_] in a clockwise manner: neurons in the quiescent state 0 spike, hence their state variable becomes 1; similarly, spiking neurons become refractory. Of the neurons in the refractory state, those being the ones nearest *ξ*_1_(*t*), a proportion *p* become quiescent, while the remaining ones remain refractory.
3. At the back of the wave, to the left of *ξ*_1_(*t*), the interval contains a mixture of neurons in states 0 and –1. The former remain at 0 whilst, of the latter, a proportion *p* transition into state 0, with the rest remaining at –1 (rules for *x* ∉ [*ξ*_1_, *ξ*_2_]). From this argument, we see that the proportion of refractory neurons in the back of the wave must decrease as *ξ* → –*π*.

**Fig. 5.**
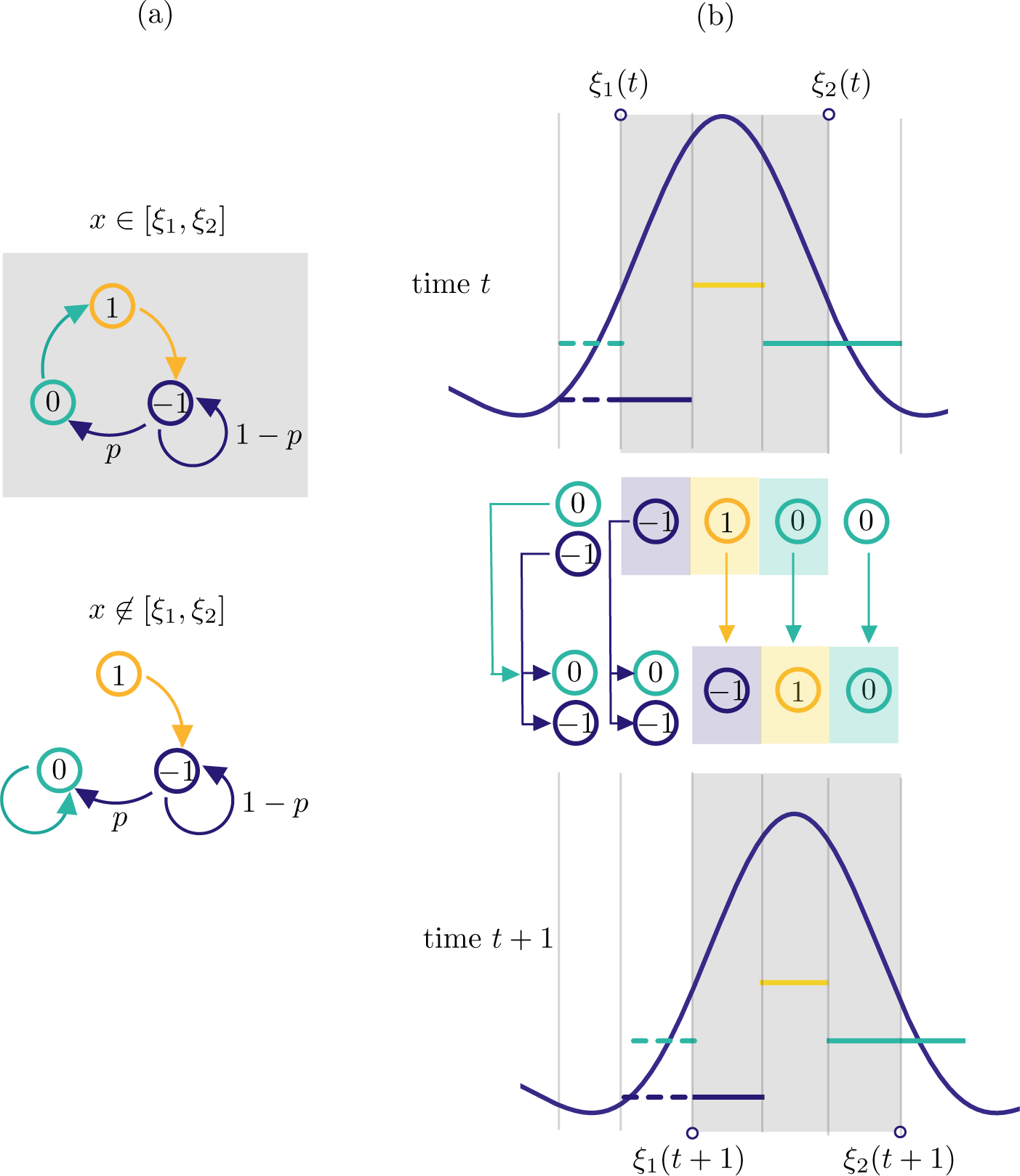
Schematic of the advection mechanism for the travelling wave state, Shaded areas pertain to the active region [*ξ*_1_(*t*), *ξ*_2_(*t*)], non-shaded areas to the inactive region 𝕏 \ [*ξ*_1_(*t*), *ξ*_2_ (*t*)]. (a): In the active (inactive) region, *q*_*i*_ = *f*(*J*(*u*))(*x*_*i*_) ≈ 1 (*q*_*i*_ ≈ 0), hence the transition kernel (2.5)–(2.7) can be simplified as shown, (b): At time *t* the travelling wave has a profile similar to the one in Figure 4, which we represent in the proximity of the active region, We depict 5 intervals of equal width, 3 of which form a partition of [*ξ*_1_(*t*), *ξ*_2_(*t*)]. Each interval is mapped to another interval at time *t* + 1, following the transition rules sketched in (a), In one discrete step, the wave progresses with positive speed: so that *J*(*x*, *t* + 1) is a translation of *J*(*x*, *t*).

The resulting mesoscopic variable *J*(*x*, *t* + 1) is a spatial translation by (*ξ*_2_(*t*) – *ξ*_1_ (*t*))/3 of *J*(*x*,*t*). We remark that the approximate transition rules of Figure 5(a) are valid also in the case of a bump, albeit the corresponding microstructure does not allow the advection mechanism described above.

### 3.4. Macroscopic variables

The computations of the previous sections suggest that, beyond the mesoscopic variable, *J*(*x*), coarser macroscopic variables are available to describe the observed patterns. In analogy with what is typically found in neural fields with Heaviside firing rate [2, 12, 18], the scalars {*ξ*_*i*_} defining the active region *X*_≥_ = ∪_*i*_ [*ξ*_2*i*–1_, *ξ*_2*i*_], where *J* is above *h*, seem plausible macroscopic variables. This is evidenced not only by Figures 2–4, but also from the schematic in Figure 5(b), where the interval [*ξ*_1_(*t*), *ξ*_2_(*t*)] is mapped to a new interval [*ξ*_1_(*t* + 1), *ξ*_2_(*t* + 1)] of the same width. To explore this further, we extract the widths Δ_*i*_(*t*) of each sub-interval [*ξ*_2*i*_(*t*), *ξ*_2*i*–1_(*t*)] from the data in Figure 2–4, and plot the widths as a function of t. In all cases, we observe a brief transient, after which Δ_*i*_(*t*) relaxes towards a coarse equilibrium, though fluctuations seem larger for the bump and multiple bump when compared with those for the wave. In the multiple bump case, we also notice that all intervals have approximately the same asymptotic width.

## 4. Deterministic model

We now introduce a deterministic version of the stochastic model considered in Section 2.2, which is suitable for carrying out analytical calculations. We make the following assumptions:

1. *Continuum neural tissue*. We consider the limit of infinitely many neurons and pose the model on 𝕊.
2. *Deterministic transitions*. We assume *p* = 1, which implies a deterministic transition from refractory states to quiescent ones (see Equation (2.5)), and *β* → ∞, which induces a Heaviside firing rate *f*(*I*) = Θ(*I* – *h*) and hence a deterministic transition from quiescent states to spiking ones given sufficiently high input (see Equations (2.4), (2.6)).

In addition to the pullback sets *X*_–1_, *X*_0_, and *X*_1_ defined in (2.2), we will partition the tissue into *active* and *inactive* regions

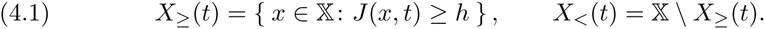

In the deterministic model, the transitions (2.5)–(2.7) are then replaced by the following rule

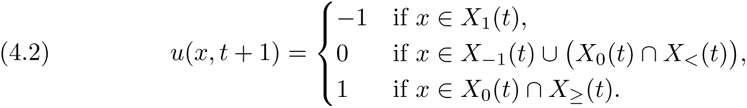

We stress that the right-hand side of the equation above depends on *u*(*x*, *t*), since the partitions {*X*_–1_, *X*_0_, *X*_1_} and {*X*_<_, *X*_≥_} do so (see Remark 2.1).

As we shall see, it is sometimes useful to refer to the induced mapping of the pullback sets

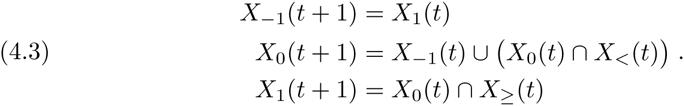

Henceforth, we will use the term *deterministic model* and formally write

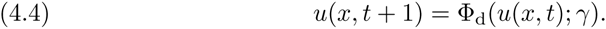

for (4.2), where the partition {*X*_*k*_}_*k*∈𝕌_ is defined by (2.2)and the active and inactive sets *X*_≥_, *X*_<_ by (4.1).

## 5. Macroscopic bump solution of the deterministic model

We now proceed to construct a bump solution of the deterministic model presented in Section 4. In order to do so, we consider a microscopic state with a regular structure, resulting in a partition, 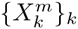, with 3*m* + 2 strips (see Figure 7) and then study the limit *m* → *∞*.

**Fig. 6.**
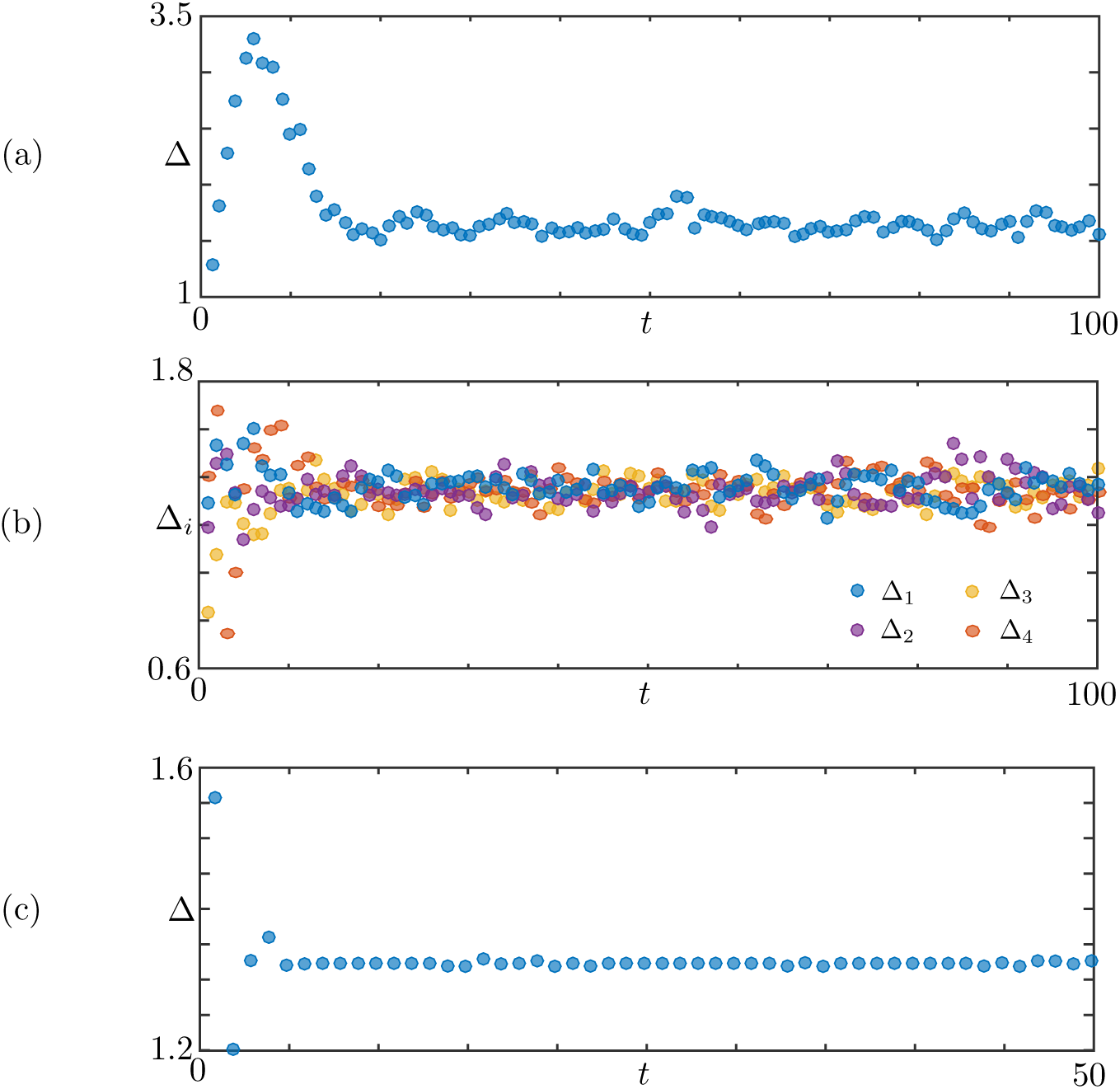
Width of the active regions Δ_*i*_ = *ξ*_2*i*_ – *ξ*_2*i*–1_ for the patterns in Figures 2–4. (a): Bump, for which *i* = 1. (b): Multiple Bump, *i* = 1,…, 4. (c): Travelling wave, *i* = 1. In all cases, the patterns reach a coarse equilibrium state after a short transient.

**Fig. 7.**
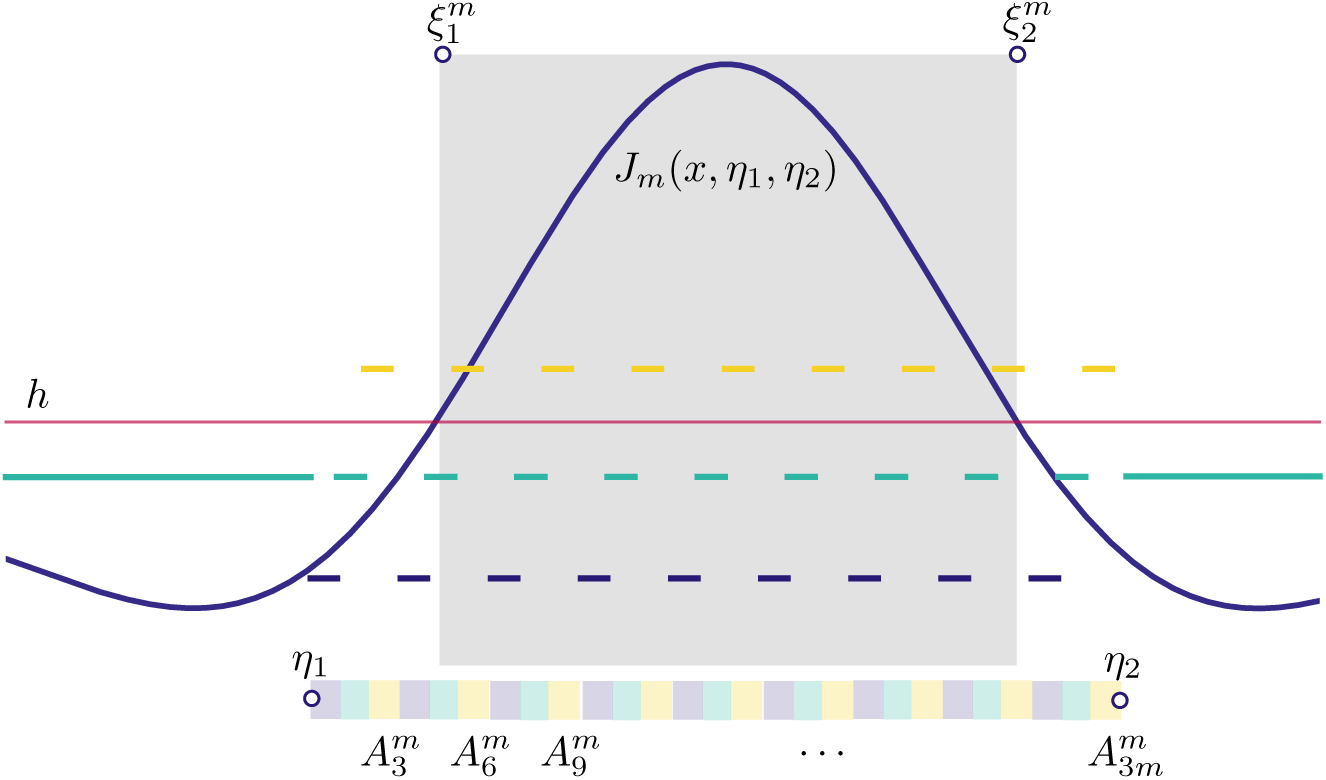
Schematic of the analytical construction of a bump, A microscopic state whose partition comprises 3m + 2 strips is considered, The microscopic state, which is not an equilibrium of the deterministic system, has a characteristic width *η*_2_ – *η*_ι_, which differs from the width 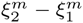 of the mesoscopic bump *J*_*m*_, If we let *m* ∞ ∞ while keeping *η*_2_ – *η*_1_ constant, then *J*_*m*_ tends towards a mesoscopic bump *J*_b_ and 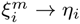 (see Proposition 5.1).

### 5.1. Bump construction

Starting from two points *η*_1_, *η*_1_ ∈ 𝕊, with *η*_1_ , *η*_2_, we construct 3*m* intervals as follows

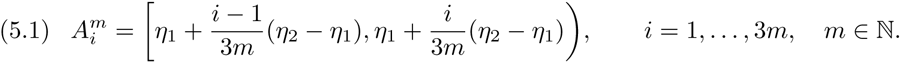

We then consider states 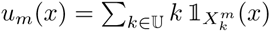, with partitions given by

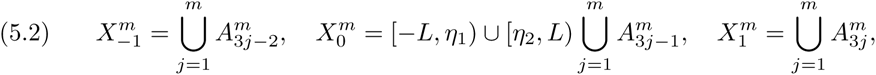

and activity set 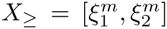. We note that, in addition to the 3m strips that form the active region of the bump, we also need two additional strips in the inactive region to form a partition of 𝕊. In general, 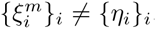, as illustrated in Figure 7. Applying (4.2)or (4.3), we find Φ_d_(*u*_*m*_) ≠ *u*_*m*_, hence *u*_*m*_ are not equilibria of the deterministic model. However, these states help us defining a macroscopic bump as a fixed point of a suitably defined map using the associated mesoscopic synaptic profile

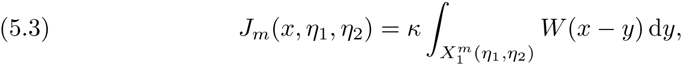

where we have highlighted the dependence of 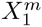 on *η*_1_, *η*_2_. The proposition below shows that there is a well defined limit, *J*_b_, of the mesoscopic profile as *m* → ∞. We also have that 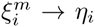 as *m* → ∞ and that the threshold crossings of the activity set are roots of a simple nonlinear function.

#### Proposition 5.1 (Bump construction).

Let *W* be the periodic extension of the synaptic kernel (2.1) and let *h*, *κ* ∈ ℝ_+_. Further, let 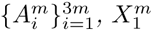 and *J*_*m*_ be defined as in (5.1), (5.2) and (5.3), respectively, and let *J*_b_: 𝕊^3^ → ℝ be defined as

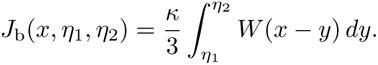

The following results hold

1. *J*_*m*_(*x*, *η*_1_, *η*_1_) → *J*_b_(*x*, *η*_1_,*η*_1_) as *m* → ∞ uniformly in the variable *x* for all *η*_1_, *η*_2_ ∈ *S* with *η*_1_ < *η*_2_,
2. If there exists Δ ∈ (0, *L*) such that 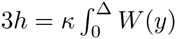, then

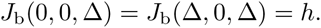

*Proof*. We fix *η*_1_ < *η*_2_ and consider the 2*L*-periodic continuous mapping *x* ↦ *J*_b_(*x*, *η*_1_, *η*_2_), defined on 𝕊. We aim to prove that *J*_*m*_ → *J*_b_ uniformly in 𝕊. We pose

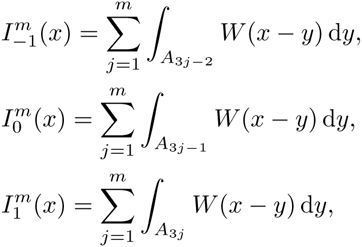

for all *x* ∈ 𝕊, *m* ∈ ℕ. Since the intervals 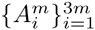 form a partition of [*η*_1_, *η*_2_) we have

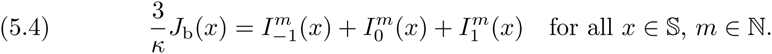

Since *W* is continuous on the compact set 𝕊, it is also uniformly continuous on 𝕊. Hence, there exists a modulus of continuity *ω* of *W*:

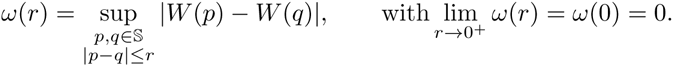

We use *ω* to estimate 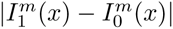 as follows:

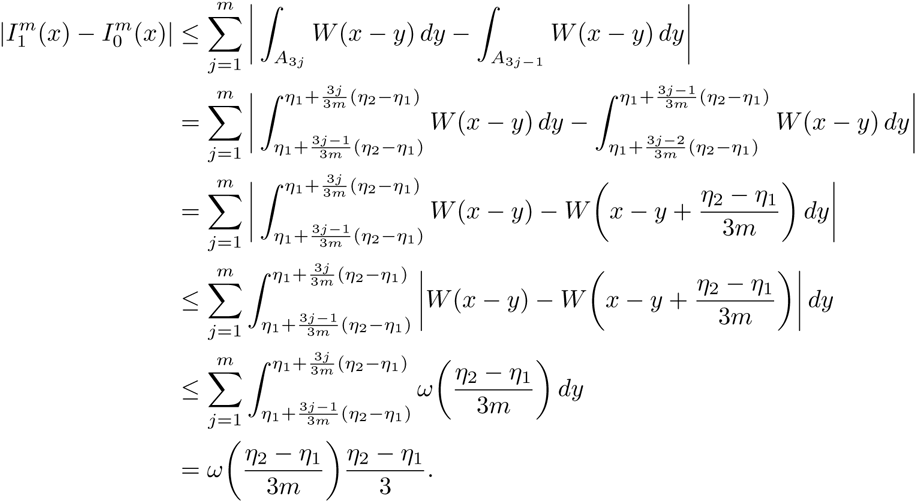

We have then 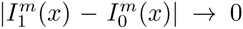 as *m* → ∞ and since *ω*((*η*_2_ – *η*_1_)/(3*m*)) is independent of *x*, the convergence is uniform. Applying a similar argument, we find 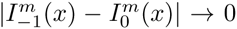 as *m* → ∞ and using (5.4), we conclude 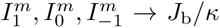 as *m* → ∞. Since 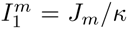, then *J*_*m*_ → *J*_b_ uniformly for all *x* ∈ 𝕊 and *η*_1_, *η*_2_ ∈ 𝕊 with *η*_1_ < *η*_2_, that is, result 1 holds true.

By hypothesis *J*_b_(0, 0, Δ) = *h* and, using a change of variables under the integral and the fact that *W* is even, it can be shown that *J*_b_(Δ, 0, Δ) = *h*, which proves result 2.

#### Corollary 5.2 (Bump symmetries).

Let Δ be defined as in Proposition 5.1, then *J*_b_(*x* + *δ*, *δ*, *δ* + Δ) is a mesoscopic bump for all *δ* ∈ [*L*, –Δ + *L*). Such bump is symmetric with respect to the axis *X* = *δ* + Δ/2.

*Proof*. The assertion is obtained using a change of variables in the integral defining *J*_b_ and noting that *W* is even.

The results above show that, 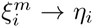 as *m* → ∞, hence we lose the distinction between width of the microscopic pattern, *η*_2_ – *η*_1_, and width of the mesoscopic pattern, 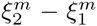, in that result 2 establishes *J*_b_(*η*_*i*_, *η*_1_, *η*_2_) = *h*, for *η*_1_ = 0, *η*_2_ = Δ. With reference to Figure 7, the factor 1/3 appearing in the expression for *J*_b_ confirms that, in the limit of infinitely many strips, only a third of the intervals 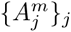 contribute to the integral. In addition, the formula for J_b_ is useful for practical computations as it allows us to determine the width, Δ, of the bump.

#### Remark 5.3 (Permuting intervals 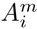).

A bump can also be found if the partition 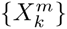 of *u*_*m*_ is less regular than the one depicted in Figure 7. In particular, Proposition 5.1 can be extended to a more general case of permuted intervals. More precisely, if we consider permutations, *σ*_*j*_, of the index sets {3*j* – 2, 3*j* – 1, 3*j*} for *J* = 1,…, *m* and construct partitions

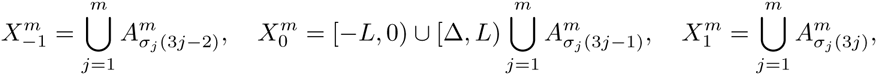

then the resulting *J*_*m*_ converges uniformly to *J*_b_ as *m* → ∞. The proof of this result follows closely the one of Proposition 5.1 and is omitted here for simplicity.

### 5.2. Bump stability

Once a bump has been constructed, its stability can be studied by employing standard techniques used to analyse neural field models [11]. We consider the map

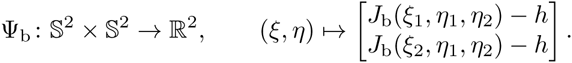

and study the implicit evolution

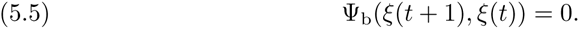

The motivation for studying this evolution comes from Proposition 5.1, according to which the macroscopic bump *ξ*_*_ = (0, Δ) is an equilibrium of (5.5), that is, Ψ_b_(*ξ*_*_, *ξ*_*_) = 0. To determine coarse linear stability, we study how small perturbations of *ξ*_*_ evolve according to the implicit rule (5.5). We set *ξ*(*t*) = *ξ*_*_ + ε*ξ*̃(*t*), for 0 < *ϵ* ≪ 1 with *ξ*̃_*i*_ = ***O***(1) and expand (5.5) around (*ξ*_*_, *ξ*_*_), retaining terms up to order *ε*,

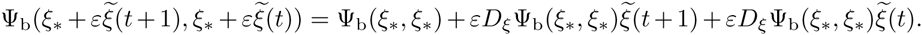

Using the classical ansatz *ξ*̃(*t*) = λ^*t*^*v*, with λ ∈ ℂ and *v* ∈ 𝕊^2^, we obtain the eigenvalue problem

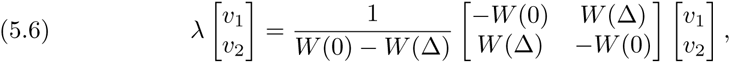

with eigenvalues and eigenvectors given by

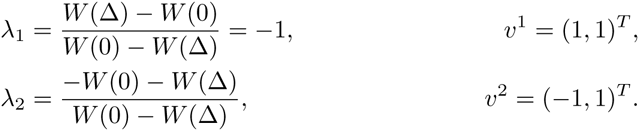

As expected, we find an eigenvalue with absolute value equal to 1, corresponding to a pure translational eigenvector. The remaining eigenvalue, corresponding to a compression/extension eigenvector, determines the stability of the macroscopic bump. The parameters *A*_*i*_, *B*_*i*_ in Equation (2.1) are such that *W* has a global maximum at *x* = 0, with *W*(0) > 0. Hence, the eigenvalues are finite real numbers and the pattern is stable if *W*(Δ) < 0. We will present concrete bump computations in Section 10.

### 5.3. Multi-bump solutions

The discussion in the previous section can be extended to the case of solutions featuring multiple bumps. For simplicity, we will discuss here solutions with 2 bumps, but the case of k bumps follows straightforwardly. The starting point is a microscopic structure similar to (5.2), with two disjoint intervals [*η*_1_, *η*_2_), [*η*_3_, *η*_4_) ⊂ 𝕊 each subdivided into 3*m* subintervals. We form the vector 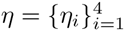 and have

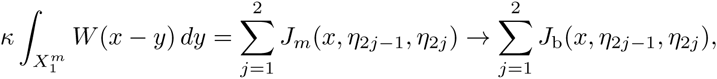

as *m* → ∞ uniformly in the variable *x* for all *η*_1_,…, *η*_4_ ∈ 𝕊 with *η*_1_ < … < *η*_4_. In the expression above, *J*_*m*_ and *J*_b_ are the same functions used in Section 5.1 for the single bump. In analogy with what was done for the single bump, we consider the mapping defined by

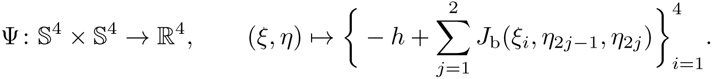

Multi-bump solutions can then be studied as in Section 5. We present here the results for a multi-bump for *L* = *π* with threshold crossings given by

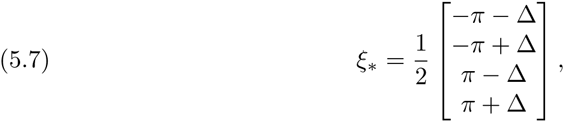

where Δ satisfies

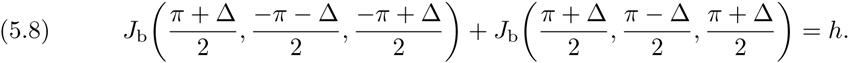

A quick calculation leads to the eigenvalue problem

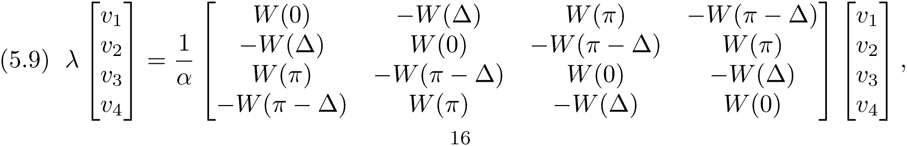

where *α* = –*W*(0) + *W*(Δ) – *W*(*π*) + *W*(*π* – Δ). The real symmetric matrix in Equation (5.9)has eigenvalues and eigenvectors given by

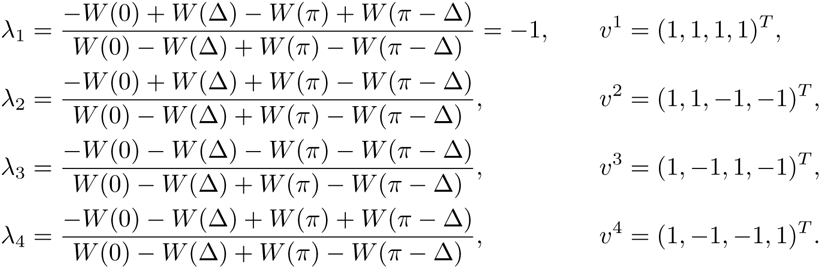

As expected, we have one neutral translational mode. If the remaining 3 eigenvalues lie in the unit circle, the multi-bump solution is stable. A depiction of this multi-bump, with corresponding eigenmodes can be found in Figure 8. We remark that the multi-bump presented here was constructed imposing particular symmetries (the pattern is even; bumps all have the same widths). The system may in principle support more generic bumps, but their construction and stability analysis can be carried out in a similar fashion.

**Fig. 8.**
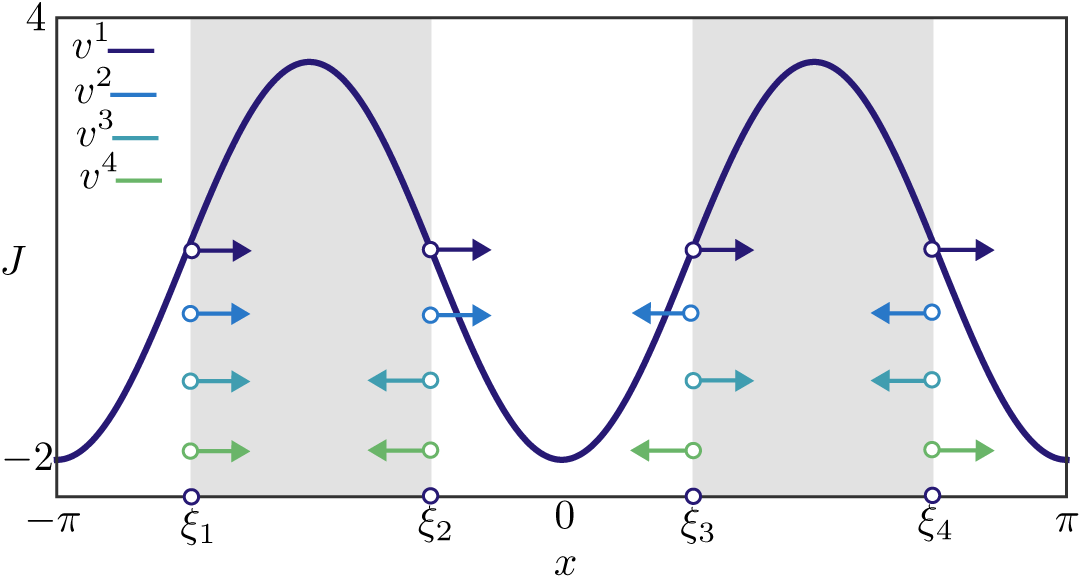
Stable mesoscopic multi-bump obtained for the deterministic model. We also plot the corresponding macroscopic bump *ξ*_*_ (Equations (5.7)-(5.8)) and coarse eigenvectors. Parameters are *k* = 30, *h* = 0.9, *p* = 1, *β* → ∞, with other parameters as in Table 1

## 6. Travelling waves in the deterministic model

Travelling waves in the deterministic model can also be studied via threshold crossings, and we perform this study in the present section. We seek a measurable function *u*_tw_ : 𝕊 → 𝕌 and a constant *c* ∈ ℝ such that

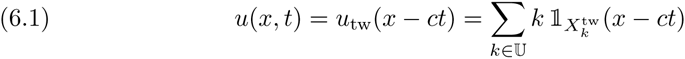

almost everywhere in 𝕊 and for all *t* ∈ ℤ. We recall that, in general, a state *u*(*x*, *t*) is completely defined by its partition, 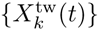. Consequently, Equation (6.1)expresses that a travelling wave has a fixed profile *u*_tw_, whose partition, 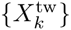, does not depend on time. A travelling wave (*u*_tw_, *c*) satisfies almost everywhere the condition

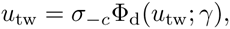

where Φ_d_ is the deterministic evolution operator (4.4)and the shift operator is defined by *σ*_*x*_ : *u*(·) ↦ *u*(· – *x*). The existence of a travelling wave is now an immediate consequence of the symmetries of *W*, as shown in the following proposition. An important difference with respect to the bump is that analytical expressions can be found for both microscopic and mesoscopic profiles, as opposed to Proposition 5.1, which concerns only the mesoscopic profile.

### Proposition 6.1 (Travelling wave)

Let *h*, *k* ∈ ℝ_+_. If there exists Δ ∈ (0, *L*) such that 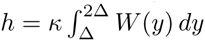, then

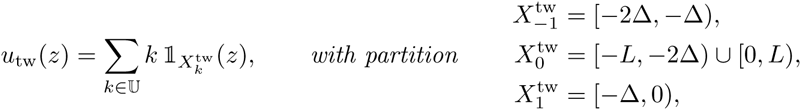

is a travelling wave of the deterministic model (4.4) with speed *c* = Δ, associated mesoscopic profile 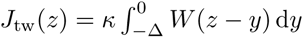 and activity set 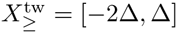.

*Proof*. The assertion can be verified directly. We have

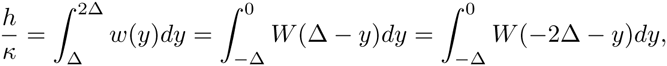

hence the activity set for *u*_tw_ is 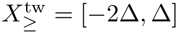 with mesoscopic profile 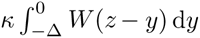. Consequently, Φ_d_(*u*_tw_; *γ*) has partition

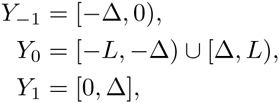

and *u*_tw_ = *σ*_–Δ_Φ_d_(*u*_tw_, *γ*) almost everywhere.

Numerical simulations of the deterministic model confirm the existence of the mesoscopic travelling wave *u*_tw_ in a suitable region of parameter space, as will be shown in Section 10. The main difference between *u*_tw_ and the stochastic waves observed in Figure 4 is in the wake of the wave, where the former features quiescent neurons and the latter a mixture of quiescent and refractory neurons.

### 6.1. Travelling wave stability

As we will show in Section 10, waves can be found for sufficiently large values of the gain parameter *κ*. However, when this parameter is below a critical value, we observe that waves destabilise at their tail. This type of instability is presented in the numerical experiment of Figure 9. Here, we iterate the dynamical system

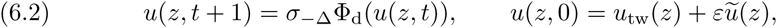

where *u*_tw_ is the profile of Proposition 6.1, travelling with speed Δ, and the perturbation *εu*̃_tw_ is non-zero only in two intervals of width 0.01. We deem the travelling wave stable if *u*(*z*, *t*) → *u*_tw_(*z*) as *t* → ∞. For *κ* sufficiently large, the perturbations decay, as witnessed by their decreasing width in Figure 9(a). For *κ* = 33, the perturbations grow and the wave destabilises.

**Fig. 9.**
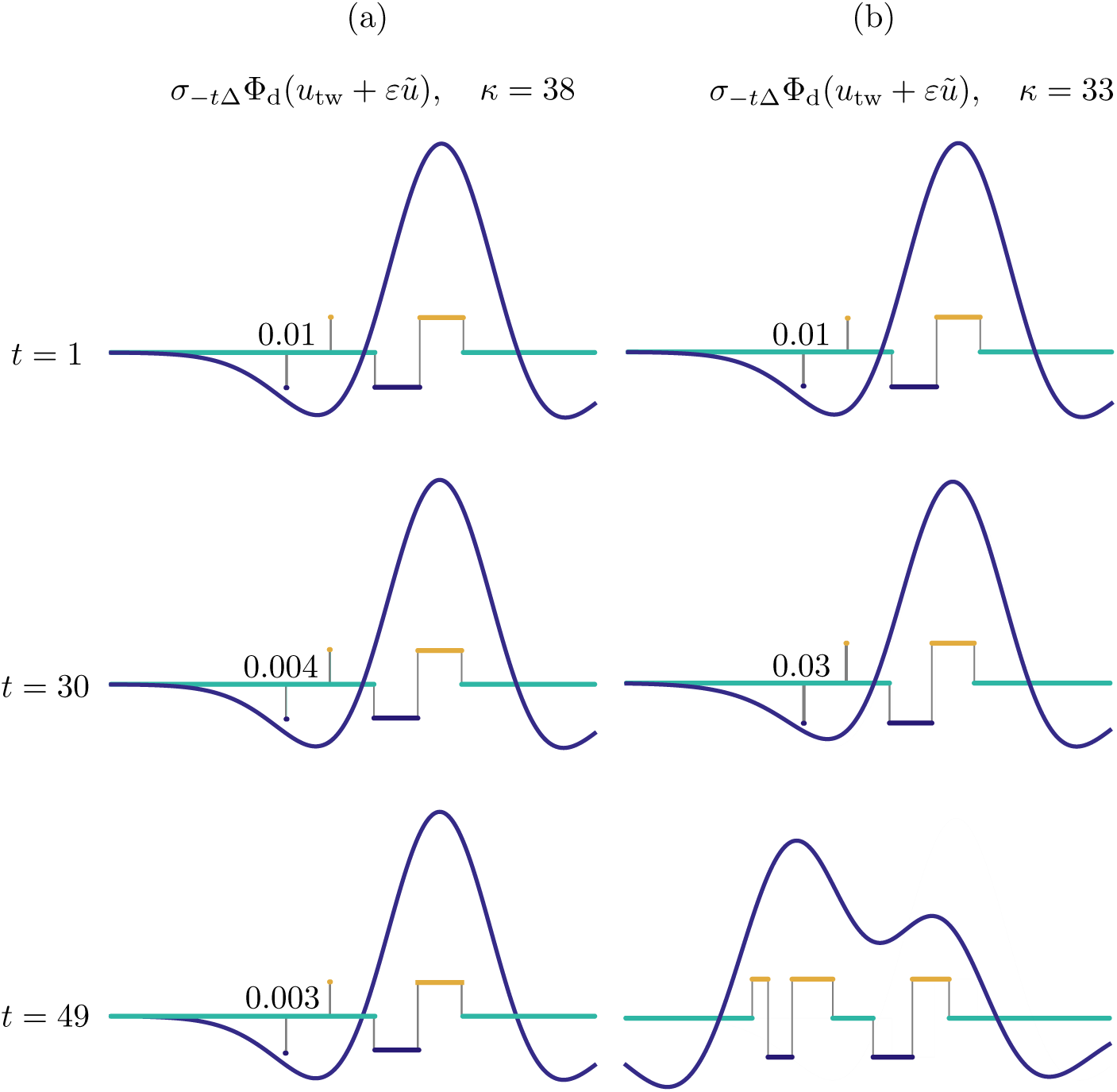
Numerical investigation of the linear stability of the travelling wave of the deterministic system, subject to perturbations in the wake of the wave. We iterate the map Φ_d_ starting from a perturbed state *u*_tw_ + *εu*̃, where *u*_tw_ is the mesoscopic wave profile of Proposition 6.1, travelling with speed Δ, and ε*u*̃ is non-zero only in two intervals of width 0.01 in the wake of the wave. We plot *σ*_–*t*Δ_Φ_d_(*u*_tw_ + *ε*_*u*̃_) and the corresponding macroscopic profile as a function of *t* and we annotate the width of one of the perturbations. (a): For *k* = 38, the wave is stable. (b): for sufficiently small *k*, the wave becomes unstable.

To analyse the behaviour of Figure 9, we shall derive the evolution equation for a relevant class of perturbations to *u*_tw_. This class may be regarded as a generalisation of the perturbation applied in this figure and is sufficient to capture the instabilities observed in numerical simulations. We seek solutions to (6.2) with initial condition *u*(*z*, *t*) = Σ_*k*_*k***1***X*_*k*(*t*)_(*z*) with time-dependent partitions

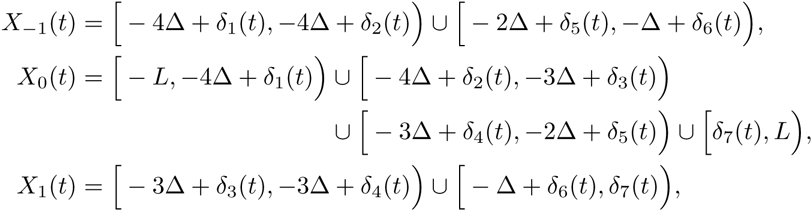

and activity set *X*_≥_(*t*) = [*ξ*_1_(*t*), *ξ*_2_(*t*)]. In passing, we note that for *δ*_*i*_ = 0, the partition above coincides with 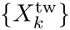 in Proposition 6.1, hence this partition can be used as perturbation of *u*_tw_. Inserting the ansatz for *u*(*ξ*, *t*) into (6.2), we obtain a nonlinear implicit evolution equation, Ψ(*δ*(*t* + 1), *δ*(*t*)) = 0, for the vector *δ*(*t*) as follows (see Figure 10)

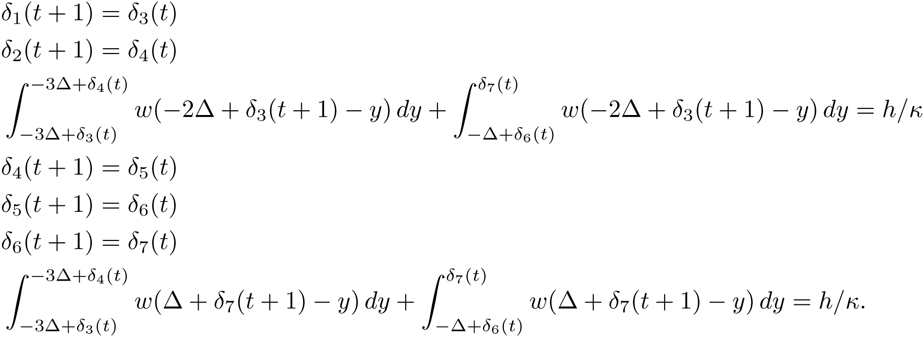

We note that the map above is valid under the assumption *δ*_3_(*t*) < *δ*_4_(*t*), which preserve the number of intervals of the original partition. As in [44], we note that this prevents us from looking at oscillatory evolution of *δ*(*t*). We set *δ*_*i*_(*t*) = *ελ*^*t*^*v*_*i*_, retain terms up to first order and obtain an eigenvalue problem for the matrix

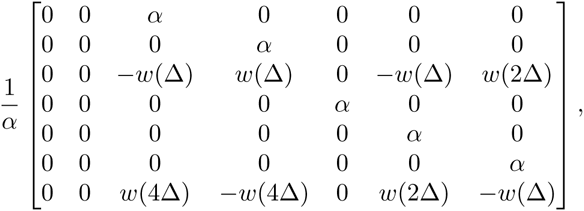

where *α* = *ω*(2Δ) – *ω*(Δ). Once again, we have an eigenvalue on the unit circle, corresponding to a neutrally stable translation mode. If all other eigenvalues are within the unit circle, then the wave is linearly stable. Concrete calculations will be presented in Section 10.

**Fig. 10.**
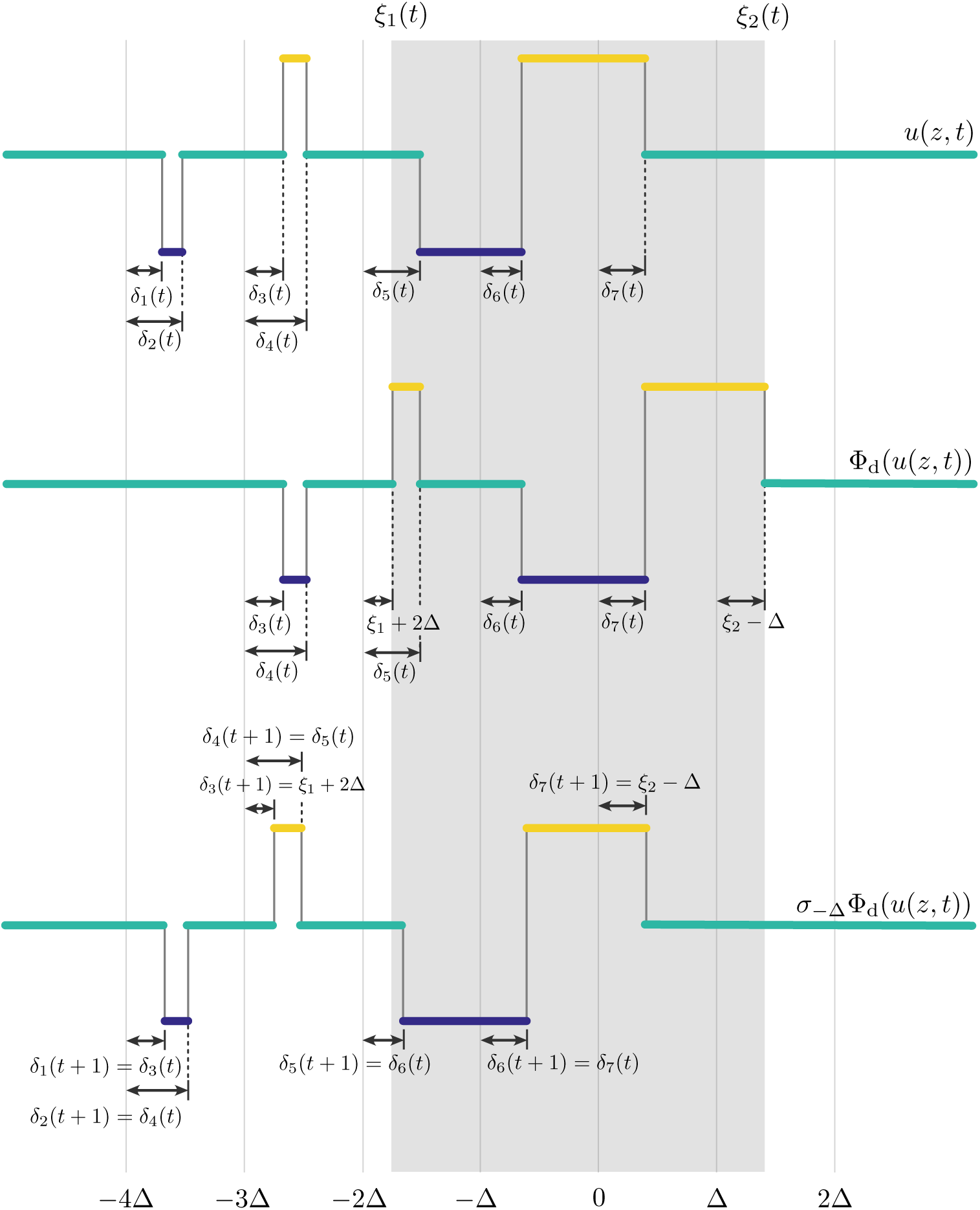
Visualisation of one iteration of the system (6.2): a perturbed travelling wave (top) is first transformed by Φ_d_ using the rules (4.3) (centre) and then shifted back by an amount Δ (bottom), This gives rise to an implicit evolution equation Ψ(*δ*(*t* + 1), *δ*(*t*)) = 0 for the threshold crossing points of the perturbed wave, as detailed in the text.

## Approximate probability mass functions for the Markov chain model

We have thus far analysed coherent states of a deterministic limit of the Markov chain model, and we now move to the more challenging stochastic setting. More precisely, we return to the original model (2.8)and find *approximate* mass functions for the coherent structures presented in Section 3 (see Figures 2–4). These approximations will be used in the lifting procedure of the equation-free framework.

The stochastic model is a Markov chain whose 3^*N*^-by-3^*N*^ transition kernel has entries specified by (2.1). It is useful to examine the evolution of the probability mass function for the state of a neuron at position *x*_*i*_ in the network, *μ*_*k*_(*x*_*i*_, *t*) = Pr(*u*(*x*_*i*_, *t*) = *k*), *k* ∈ 𝕌, which evolves according to

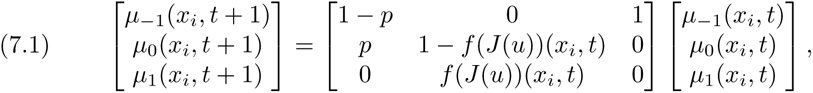

or in compact notation *μ*(*x*_*i*_, *t* + 1) = Π(*x*_*i*_, *t*)*μ*(*x*_*i*_, *t*). We recall that *f* is the sigmoidal firing rate and that *J* is a deterministic function of the random vector, *u*(*x*, *t*) ∈ 𝕌^*N*^, via the pullback set 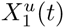:

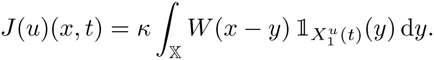

As a consequence, the evolution equation for *μ*(*x*_*i*_, *t*) is non-local, in that *J*(*x*_*i*_, *t*) depends on the microscopic state of the whole network.

We now introduce an approximate evolution equation, obtained by posing the problem on a continuum tissue 𝕊 and by substituting *J*(*x*, *t*) by its expected value

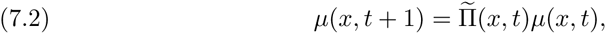

where *μ* : 𝕊 × ℤ → [0,1]^3^,

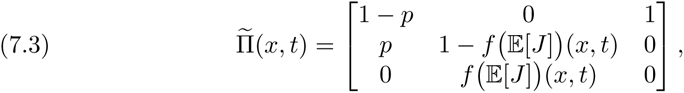

and

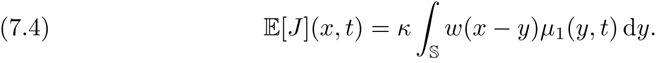

In passing, we note that the evolution equation (7.2)is deterministic. We are interested in two types of solutions to (7.2):

1. A time-independent bump solution, that is a mapping *μ*_b_ such that *μ*(*x*, *t*) = *μ*_b_(*x*) for all *x* ∈ 𝕊 and *t* ∈ ℤ.
2. A travelling wave solution, that is, a mapping *μ*_tw_ and a real number *c* such that *μ*(*x*, *t*) = *μ*_tw_(*x* – *ct*) for all *x* ∈ 𝕊 and *t* ∈ ℤ.

### 7.1. Approximate probability mass function for bumps

We observe that, posing *μ*(*y*, *t*) = *μ*_b_(*y*) in (7.2), we have

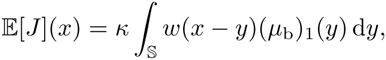

Motivated by the simulations in Section 3 and by Proposition 5.1, we seek a solution to (7.2) in the limit *β* → œ, with 𝔼[*J*](*x*) ≥ *h* for *x* ∈ [0, Δ], and (*μ*_b_)_1_(*x*) ≠ 0 for *x* ∈ [0, Δ], where Δ is unknown. We obtain

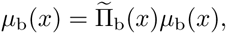

where

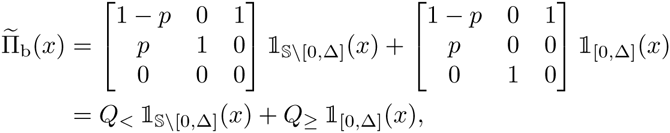

We conclude that, for each *x* ∈ [0, Δ] (respectively *x* ∈ 𝕊 \ [0, Δ]), *μ*_b_(*x*) is the right || · ||_1_-unit eigenvector corresponding to the eigenvalue 1 of the stochastic matrix *Q*_≥_ (respectively *Q*_<_). We find

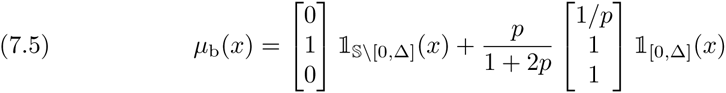

and, by imposing the threshold condition 𝔼[*J*](Δ) = *h*, we obtain a compatibility condition for Δ,

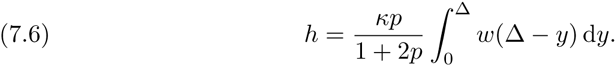

We note that if *p* = 1 we have 𝔼[*J*](*x*) = *J*_b_ (*x*, 0, Δ) where *J*_b_ is the profile for the mesoscopic bump found in Proposition 5.1, as expected.

In Figure 11(a), we plot *μ*_b_(*x*) as predicted by (7.5)–(7.6), for *p* = 0.7, *κ* = 30, *h* = 0.9. At each *x*, we visualise (*μ*_b_)_*k*_ for each *k* ∈ 𝕌 using vertically juxtaposed color bars, with height proportional to the values (*μ*_b_)_*k*_, as shown in the legend. For a qualitative comparison with direct simulations, we refer the reader to the microscopic profile *u*(*x*, 50) shown in the right panel of Figure 2(a): the comparison suggests that each *u*(*x*_*i*_, 50) is distributed according to *μ*_b_(*x*_*i*_).

**Fig. 11.**
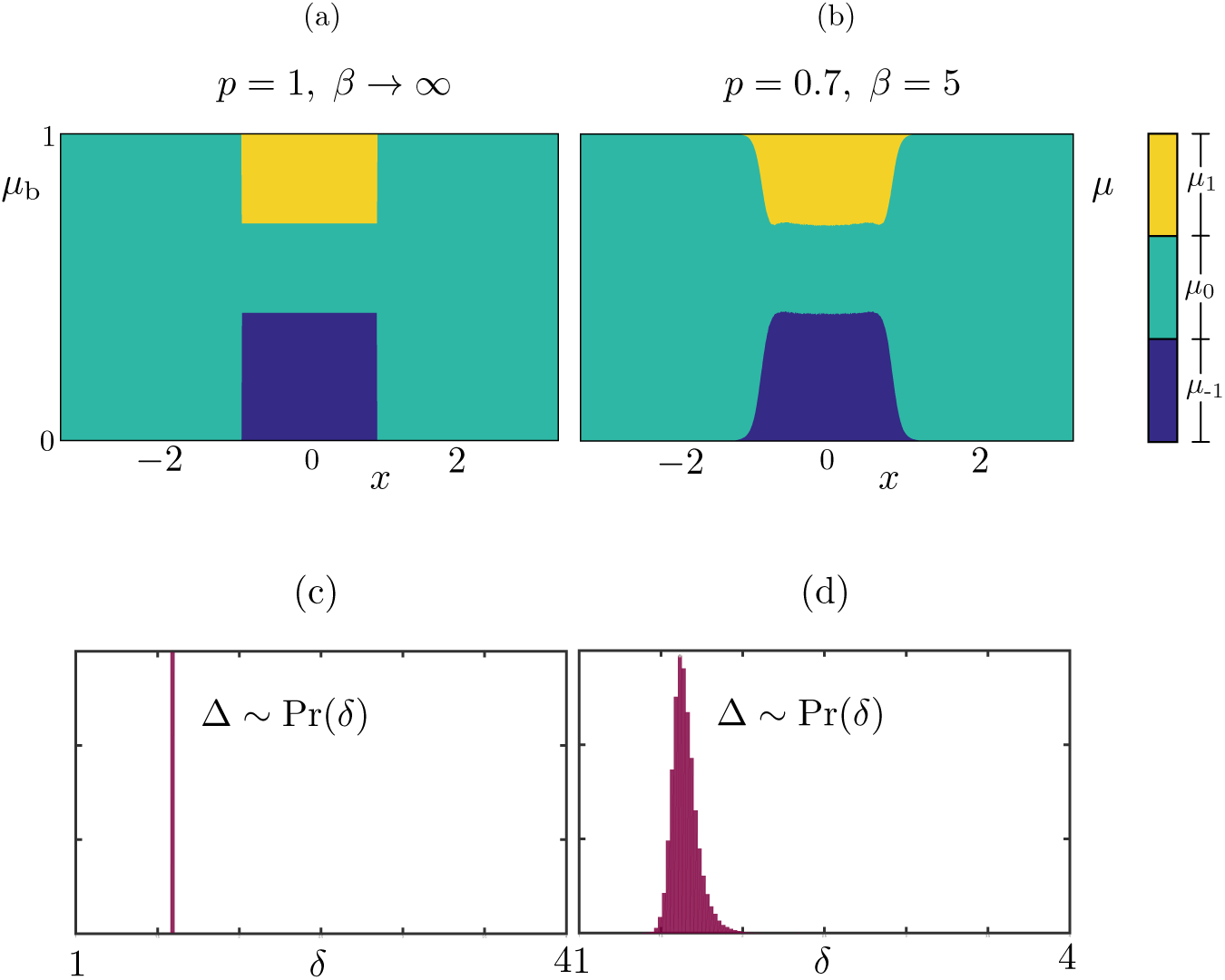
Comparison between the probability mass function μb, as computed by (7.5)-(7.6), and the observed distribution μ of the stochastic model, (a): We compute the vector (*μ*_b_)_*k*_, *k* ∈ 𝕌 in each. strip using (7.7) and visualise the distribution using vertically juxtaposed color bars, with height proportional to the values (*μ*_b_)_*k*_, as shown in the legend, (b): A long. simulation of the stochastic model supporting a stochastic bump *u*(*x*,*t*) for *t* ∈ [0,*T*], where *T* = 10^5^, At each time *t* > 10 (allowing for initial transients to decay), we compute *ξ*_1_(*t*), *ξ*_2_(*t*), Δ(*t*) and then produce histograms for the random profile *u*(*x* – *ξ*_1_(*t*) – Δ(*t*)/2,*t*), (c): in the deterministic limit the value of Δ is determined by (7.6), hence we have a Dirac distribution, (d): the distribution of Δ obtained in the Markov chain model, Parameters are as in Table 1.

We also compared quantitatively the approximate distribution *μ*_b_ with the distribution, *μ*(*x*, *t*), obtained via Monte Carlo samples of the full system (7.1). The distributions are obtained from a long-time simulation of the stochastic model supporting a microscopic bump *u*(*x*, *t*) for *t* ∈ [0,*T*], with *T* = 10^5^. At each discrete time *t*, we compute the mesoscopic profile, *J*(*u*)(*x*, *t*), the corresponding threshold crossings and width: *ξ*_1_(*t*), *ξ*_2_(*t*), Δ(*t*) and then produce histograms for the random profile *u*(*x* – *ξ*_1_(*t*) – Δ(*t*)/2, *t*). The instantaneous shift applied to the profile is necessary to pin the wandering bump.

We note a discrepancy between the analytically computed histograms, in which we observe a sharp transition between the region *x* ∈ [0, Δ] and *x* ∈ 𝕊 \ [0, Δ], and the numerically computed ones, in which this transition is smoother. This discrepancy arises because Δ(*t*) oscillates around an average value Δ predicted by (7.6); the approximate evolution equation (7.2)does not account for these oscillations. This is visible in the histograms of Figure 11(c)-(d), as well as in the direct numerical simulation 6(a).

### 7.2. Approximate probability mass function for travelling waves

We now follow a similar strategy to approximate the probability mass function for travelling waves. We pose *μ*(*x*, *t*) = *μ*_tw_(*x* – *ct*) in the expression for 𝔼[*J*], to obtain

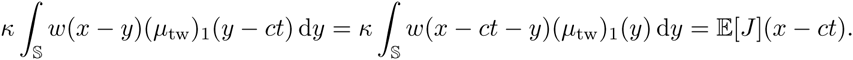

Proposition 6.1 provides us with a deterministic travelling wave with speed *c* = Δ. The parameter Δ is also connected to the mesoscopic wave profile, which has threshold crossings *ξ*_1_ = –2Δ and *ξ*_2_ = Δ. Hence, we seek for a solution to (7.2) in the limit *β* → ∞, with 𝔼[*J*](*z*) ≥ *h* for *X* ∈ [–2*c*, *c*], and (*μ*_tw_)_1_(*z*) ≠ 0 for *z* ∈ [–2*c*, *c*], where *c* is unknown. For simplicity, we pose the problem on a large domain whose size is commensurate with *c*, that is 𝕊 = *cT*/ℝ, where *T* is an even integer much greater than 1.

We obtain

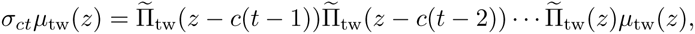

where

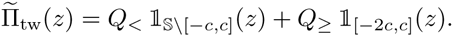

To make further analytical progress, it is useful to partition the domain 𝕊 = *cT*/ℝ in strips of width *c*,

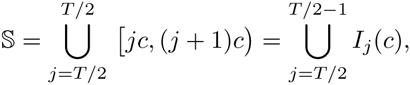

and impose that the wave returns back to its original position after *t* iterations, *σ*_*cT*_*μ*_tw_(*z*) = *μ*_tw_(*z*), while satisfying the compatibility condition *h* = 𝔼[*J*](*c*). This leads to the system

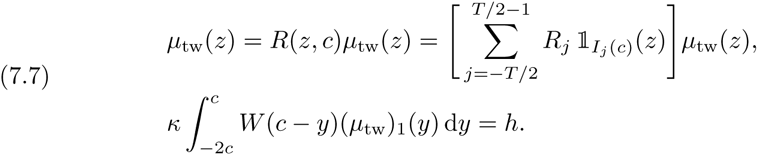

With reference to system (7.7)we note that:

1. *R*(*z*, *c*) is constant within each strip *I*_*j*_, hence the probability mass function, *μ*_tw_(*z*), is also constant in each strip, that is, *μ*_tw_(*z*) = Σ_*i*_ *ρ*_*i*_**1***I*_*i*(*c*)_(*z*) for some unknown vector (*ρ*_–*T*/2_,…, *ρ*_*T*/2_) ∈ 𝕊^3*T*^.
2. Each *R*_*j*_ is a product of *T* 3-by-3 stochastic matrices, each equal to *Q*_<_ or *Q*_≥_. Furthermore, the matrices {*R*_*j*_} are computable. For instance, for the strip *I*_–1_ we have

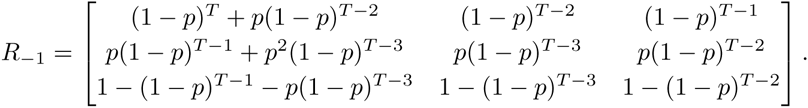

Consequently, *μ*_tw_(*z*) can be determined by solving the following problem in the unknown (*ρ*_–T/2_, …, *ρ*_*T*/2_, *c*) ∈ 𝕊^3*T*^ × ℝ:

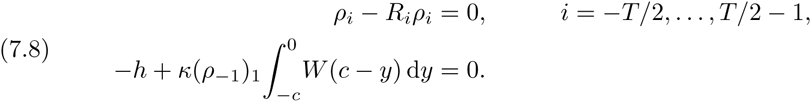

Before presenting a quantitative comparison between the numerically determined distribution, *μ*_tw_(*z*), and that obtained via direct time simulations, we make a few efficiency considerations. In the following sections, it will become apparent that sampling the distribution *μ*_tw_(*z*) for various values of control parameters, such as *h* or *κ*, is a recurrent task, at the core of the coarse bifurcation analysis: each linear and nonlinear function evaluation within the continuation algorithm requires sampling *μ*_tw_(*z*), and hence solving the large nonlinear problem (7.8).

With little effort, however, we can obtain an accurate *approximation* to *μ*_tw_, with considerable computational savings. The inspiration comes once again from the analytical wave of Proposition 6.1. We notice that only the last equation of system (7.8)is nonlinear; the last equation is also the only one which couples {*ρ*_*j*_} with *c*. When *p* = 1 the wave speed is known as *β* → ∞, *N* → ∞ and *p* = 1 corresponds to the deterministic limit, hence 𝔼[*J*](*z*) = *J*_tw_(*z*), which implies *c* = Δ and (*ρ*_–1_)_1_ = 1. The stochastic waves observed in direct simulations for *p* ≠ 1, however, display *c* ≈ (*ξ*_2_ – *ξ*_1_)/3 = Δ and μ ≈ 1 in the strip where *J* achieves a local maximum (see, for instance Figure 4, for which *p* = 0.4).

The considerations above lead us to the following scheme to approximate *μ*_tw_: (i) set *c* = Δ and remove the last equation in (7.8); (ii) solve *T* decoupled 3-by-3 eigenvalue problems to find *ρ*_*i*_. Furthermore, if *p* remains fixed in the coarse bifurcation analysis, *ρ*_*i*_ can be pre-computed and step (ii) can be skipped.

In Figure 12(a), we report the approximate *μ*_tw_ found with the numerical procedure described above. An inspection of the microscopic profile *u*(*x*, 45) in the right panel of Figure 4(a) shows that this profile is compatible with *μ*_tw_. We also compared quantitatively the approximate distribution with the distribution, *μ*(*x*, *t*), obtained with Monte Carlo samples of the full system (7.1). The distributions are obtained from *M* samples 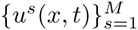 of the stochastic model for a travelling wave for *t* ∈ [0, *T*]. For each sample *s*, we compute the thresholds, 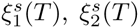, of the corresponding *J*(*u*^*s*^)(*x*, *t*) and then produce histograms for 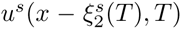. This shifting, whose results are reported in Figure 12(b), does not enforce any constant value for the velocity, hence it allows us to test the numerical approximation *μ*_tw_. The agreement between the two distributions is excellent: we stress that, while the strips in Figure 12(a) are enforced by our approximation, the ones in Figure 12(b) emerge from the data. We note a slight discrepancy, in that *μ*_tw_(– 3Δ) ≈ 0, while the other distribution shows a small nonzero probability attributed to the firing state at *ξ* = –3Δ. Despite this minor disagreement, the differences between the approximated and observed distributions remain small across all parameter regimes of note and the approximations even retain their accuracy as *β* is decreased (not shown).

**Fig. 12.**
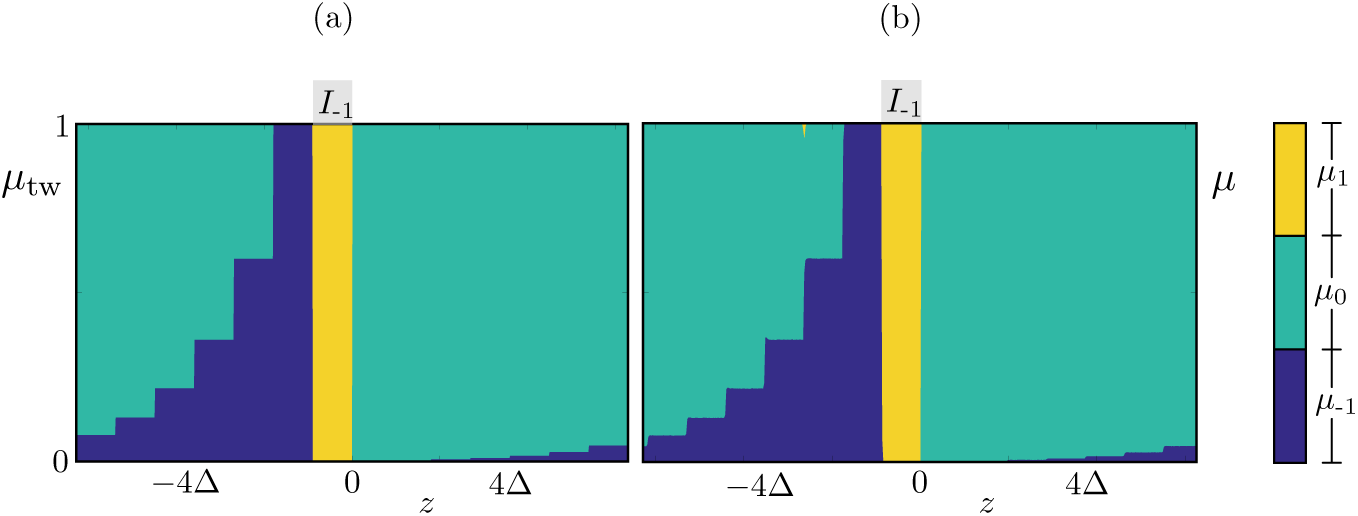
Similarly to Figure 11, we compare the approximated probability mass function *μ*, and the observed distribution μ of the stochastic model. (a): the probability mass function is approximated using the numerical scheme outlined in the main text for the solution of (7.8); the strip *I*_–1_ is indicated for reference. (b): A set of 9 × 10^5^ realisations of the stochastic model for a travelling wave are run for *t* ∈ [0,*T*], where *T* = 1000. For each realisation s, we calculate the final threshold crossings 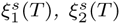, and then compute histograms of 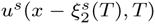. We stress that the strips in (a) are induced by our numerical procedure, while the ones in (b) emerge from the data. The agreement is excellent and is preserved across a vast region of parameter space (not shown). Parameters are as in Table 1.

## 8. Coarse time-stepper

As mentioned in the introduction, equation-free methods allow us to compute macroscopic states in cases in which a macroscopic evolution equation is not available in closed form [42, 43]. To understand the general idea behind the equation-free framework, we initially discuss an example taken from one of the previous sections, where an evolution equation *does* exist in closed form.

In Section 5, we described bumps in a deterministic limit of the Markov chain model. In this description, we singled out a *microscopic* state (the function *u*_*m*_(*x*) with partition (5.2)) and a corresponding *mesoscopic* state (the function *J*_*m*_(*x*)), both sketched in Figure 7. Proposition 5.1 shows that there exists a well defined mesoscopic limit profile, *J*_b_, which is determined (up to translations in *x*) by its threshold crossings *ξ*_1_ = 0, *ξ*_2_ = Δ. This suggests a characterisation of the bump in terms of the *macroscopic* vector (*ξ*_1_, *ξ*_2_) or, once translation invariance is factored out, in terms of the *macroscopic* bump width, Δ. Even though the microscopic state *u*_*m*_ is not an equilibrium of the deterministic system, the macroscopic state (0, Δ) is a fixed point of the evolution equation (5.5), whose evolution operator Ψ is known in closed form, owing to Proposition 5.1. It is then possible to compute Δ as a root of an explicitly available nonlinear equation.

We now aim to use equation-free methods to compute macroscopic equilibria in cases where we do not have an explicit evolution equation, but only a numerical procedure to approximate Ψ. As mentioned in the introduction, the evolution equation is approximated using a coarse time-stepper, which maps the macroscopic state at time t0 to the macroscopic state at time t1 using three stages: lifting, evolution, restriction. The specification of these stages (the lifting in particular) typically requires some closure assumptions, which are enforced numerically. In our case, we use the analysis of the previous sections for this purpose. In the following section, we discuss the coarse time-stepper for bumps and travelling waves. The multi-bump case is a straightforward extension of the single bump case.

### 8.1. Coarse time-stepper for bumps

The macroscopic variables for the bump are the threshold crossings {*ξ*_*i*_} of the mesoscopic profile *J*. The lifting operator for the bump takes as arguments {*ξ*_*i*_} and returns a set of microscopic profiles compatible with these threshold crossings:

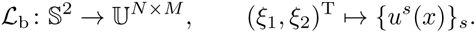

If *β* → ∞, *u*^*s*^(*x*) are samples of the analytical probability mass function *μ*_b_(*x* + Δ/2), where *μ*_b_ is given by (7.5)with Δ = *ξ*_2_ – *ξ*_1_. In this limit, a solution branch may also be traced by plotting (7.6).

If *β* is finite, we either extract samples from the *approximate* probability mass function *μ*_b_ used above, or we extract samples *U*^*s*^(*x*) satisfying the following properties (see Proposition 5.1 and Remark 5.3):

1. *U*^*s*^(*x*) is symmetric with respect to the axis *X* = (*ξ*̃_1_ + *ξ*̃_2_)/2, where *ξ*̃_*i*_ = round (*ξ*_*i*_) and round: 𝕊 → 𝕊_*N*_.
2. *u*^*s*^(*x*) = 0 for all *X* ∈ [–*L*, *ξ*̃_1_) ∪ (*ξ*̃_2_, *L*).
3. The pullback sets, *X*_1_ and *X*_–1_, are contained within [*ξ*_1_, *ξ*_2_] and are unions of a random number of intervals whose widths are also random. A schematic of the lifting operator for bumps is shown in Figure 13.

**Fig. 13.**
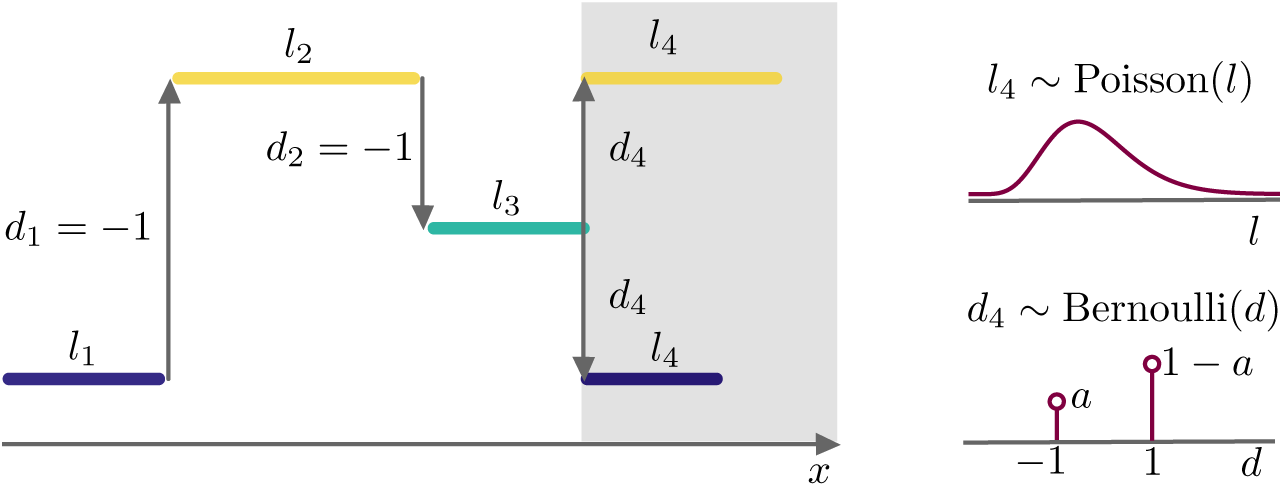
Schematic representation of the lift operator for a bump solution, This figure displays a representation of how the states for neurons located within the activity set, [*ξ*_1_,*ξ*_2_], are lifted, For illustrative purposes, we assume here that we are midway through the lifting operation, where 3 steps of the while loop listed in Algorithm 1 have been completed and a fourth one is being executed (shaded area), The width *l*_4_ of the next strip is drawn from a Poisson distribution, The random variable *d* € {–1, 1} indicates the direction through which we cycle through the states {–1, 0, 1} during the lifting, The number *d*_4_ is drawn from a Bernoulli distribution whose average a gives the probablity of changing direction, For full details of the lifting operator, please refer to Algorithm 1.

A more precise description of the latter sampling is given in Algorithm 1. As mentioned in the introduction, lifting operators are not unique and we have given above two possible examples of lifting. In our computations, we favour the second sampling method. The mesoscopic profiles, *J*, generated using this approach are well-matched to 𝔼[*J*] produced by the analytically derived probability mass functions (7.5). Numerical experiments demonstrate that this method is better than the first possible lifting choice at continuing unstable branches. This is most likely due to the fact that the latter method slightly overestimates the probability of neurons within the bump to be in the spiking state, and underestimates that of them being in the refractory state and this helps mitigate the problems encountered when finding unstable states caused by the combination of the finite size of the network and non-smooth characteristics of the model (when *β* is high).

The evolution operator is given by

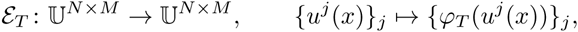

where *φ*_*T*_ denotes *T* compositions of the microscopic evolution operator (2.8)and the dependence on the control parameter, γ, is omitted for simplicity.

For the restriction operator, we compute the average activity set of the profiles. More specifically, we set

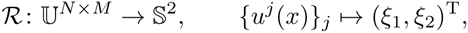

where

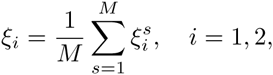

and 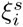 are defined using a piecewise first-order interpolant 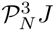 of *J* with nodes 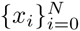,

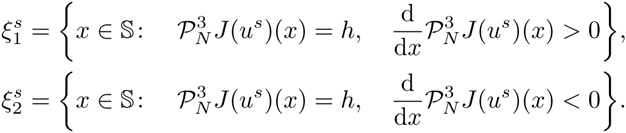

#### Algorithm 1: Lifting operator for bump

~~~
**Input**    : Threshold crossings *ξ*_1_, *ξ*_2_; Average number of strips, *m*; Average for the Bernoulli distribution, *a*; Width of the domain, *L*.
**Output**   : Profiles *u*^1^(*x*),…,*u*^*M*^(*x*)
**Comments** : The profiles *u*^*s*^(*x*) are assumed to be symmetric around x = (*ξ*_1_ + *ξ*_2_)/2. The operator *round* rounds a real number to a computational grid with stepsize *δx* = 2*L*/*N*.
**Pseudocode**      : for *s* = 1, *M* **do**
                             Set *u*^*s*^(*x*) = 0 for all *x* ∈ [–*L*, *ξ*_1_) ∪ (*ξ*_2_, *L*)
                             Set *d* = –1
                     *x* = round(*ξ*_1_), *u*^*s*^(*x*) = 1
                       **while** *x* ≤ (*ξ*_1_ + *ξ*_2_)/2 do
                                Select random width *l* ~ Poisson((*ξ*_2_ – *ξ*_1_/*m*)
                                Select random increment *b* ~ Bernoulli(*a*), *d* = (*d* + 2*b* + 1) mod 3 – 1
                          **for** *j* = 1, *l* **do**
                                   Update *x* = *x* + *δx*
                             **if** *x* ≤ (*ξ*_1_ + *ξ*_2_)/2 then
                                   **if** *j* = *l* then *u*^*i*^ changes value at the next grid point
                                    *u*^*s*^(*x* + *δx*) = (*u*^*s*^(*x*) + *d* + 1) mod 3 – 1
                                   **else** *u*^*s*^ remains constant at the next grid point
                                 *u*^*s*^(*x* + *δx*) = *u*^*s*^(*x*)
                                   **end**
                                         Reflection around symmetry axis, *u*(*ξ*_2_ + *ξ*_1_ – *x*) = *u*(*x*)
                              **end**
                        **end**
                    **end**
           **end**
~~~

We also point out that the computation stops if the two sets above are empty, whereupon, we set 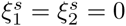.

The coarse time-stepper for bumps is then given by

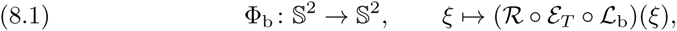

where the dependence on parameter *γ* has been omitted.

### 8.2. Coarse time-stepper for travelling waves

In Section 7.1, we showed that the probability mass function, *μ*_tw_(*z*), of a coarse travelling wave can be approximated numerically using the travelling wave of the deterministic model, by solving a simple set of eigenvalue problems. It is therefore natural to use *μ*_tw_ in the lifting procedure for the travelling wave. In analogy with what was done for the bump, our coarse variables (*ξ*_1_, *ξ*_2_) are the boundaries of the activity set associated with the coarse wave, *X*_≥_ = [*ξ*_1_, *ξ*_2_]. We then set

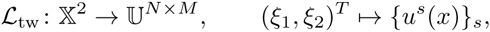

where {*u*^*s*^(*x*_*i*_)}_*s*_ are *M* independent samples of the probability mass functions *μ*_tw_(*x*_*i*_), with *c* = (*ξ*_2_ – *ξ*_1_)/3. The restriction operator for travelling waves is the same used for the bump. The coarse time-stepper for travelling waves, Φ_tw_, is then obtained as in (8.1), with 𝓛_b_ replaced by 𝓛_tw_.

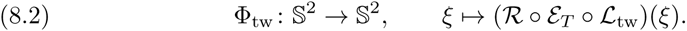

## 9. Root finding and pseudo-arclength continuation

Once the coarse time-steppers, Φ_1_ and Φ_tw_, have been defined, it is possible to use Newton’s method and pseudo-arclength continuation to compute coarse states, continue them in one of the control parameters and assess their coarse linear stability. In this section, we will indicate dependence upon a single parameter *γ* ∈ ℝ, implying that this can be any of the control parameters in (2.8).

For bumps, we continue in *γ* the nonlinear problem *F*_b_(*ξ*; *γ*) = 0, where

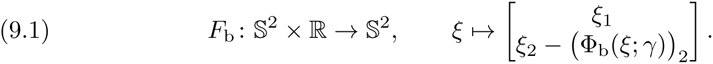

A vector *ξ* such that *F*_b_(*ξ*; *γ*) = 0 corresponds to a coarse bump with activity set *X*_≥_ = [0, *ξ*_2_] and width *ξ*_2_, occurring for the parameter value *γ*, that is, we eliminated the translation invariance associated with the problem by imposing *ξ*_1_ = 0. In passing, we note that it is possible to hardwire the condition *ξ*_1_ =0 directly in *F*_b_ and proceed to solve an equivalent 1-dimensional system. Here, we retain the 2-dimensional formulation with the explicit condition *ξ*_1_ = 0, as this makes the exposition simpler.

During continuation, the explicitly unavailable Jacobians

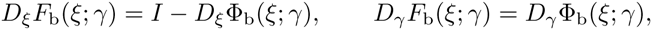

are approximated using the first-order forward finite-difference formulas

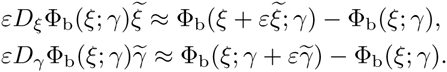

The finite difference formula for *Dξ*Φ_b_ also defines the Jacobian operator used to compute stability: for a given solution *ξ*_*_ of (9.1), we study the associated eigenvalue problem

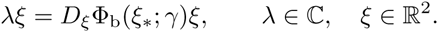

For coarse travelling waves, we define

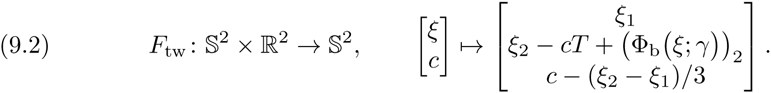

A solution (*ξ*, *c*) to the problem *F*_tw_(*ξ*, *c*; *γ*) = 0 corresponds to a coarse travelling wave with activity set *X*_≥_ = [0, *ξ*_2_] and speed *ξ*_2_/3, that is, we eliminated the translation invariance and imposed a speed c in accordance with the lifting procedure ***L***_tw_. As for the bump we can, in principle, solve an equivalent 1-dimensional coarse problem.

## 10. Numerical results

We begin by testing the numerical properties of the coarse time-stepper, the Jacobian-vector products and the Newton solver used for our computations. In Figure 14(a), we evaluate the Jacobian-vector product of the coarse time stepper with *p* = 1, *β* → ∞ for bumps (waves) evaluated at a coarse bump (wave), in the direction *εξ*̃, where 0 < *ε* ≪ 1 and *ξ*̃ is a random vector with norm 1. Since this coarse time stepper corresponds to the deterministic case, we expect the norm of the Jacobian-vector product to be an *O*(*ε*), as confirmed by the numerical experiment. In Figure 14(b), we repeat the experiment in the stochastic setting (*p* = 0.4), for the travelling wave case with various number of realisations. As expected, the norm of the Jacobian-vector action follows the *O*(*ε*) curve for sufficiently large *ε*: the more realisations are employed, the more accurately the *O*(*ε*) curve is followed.

**Fig. 14.**
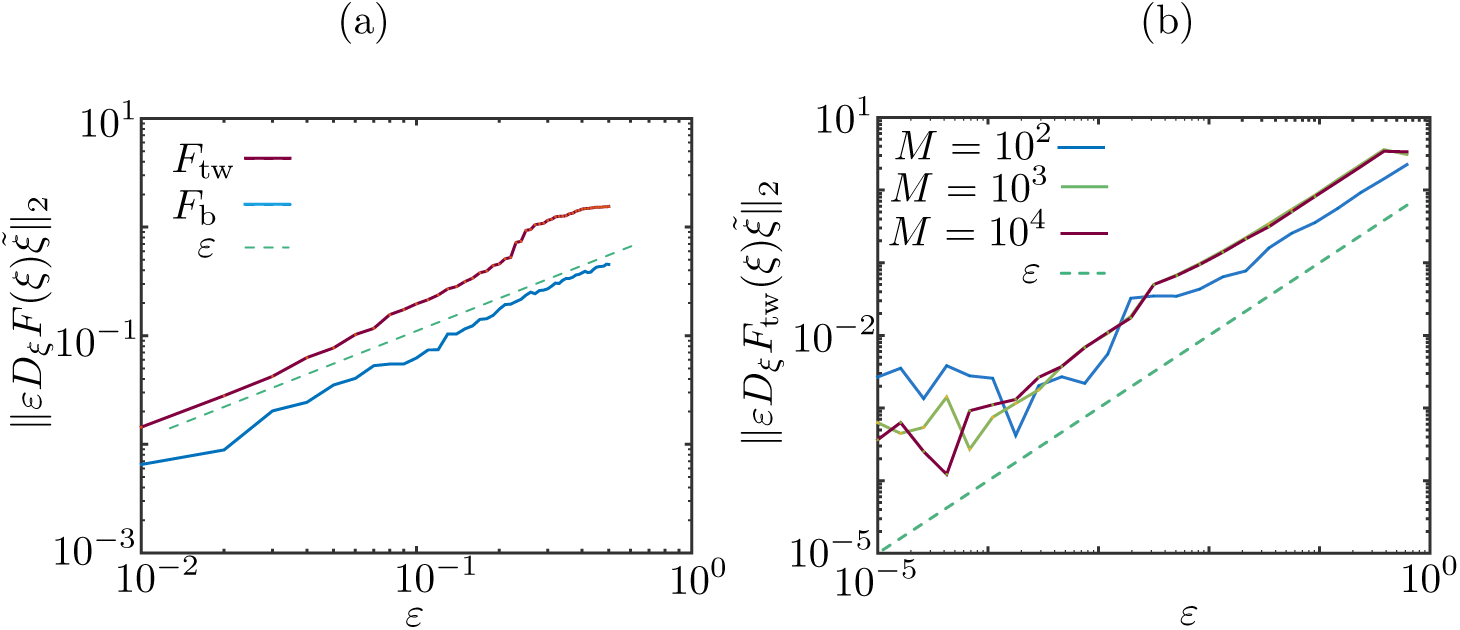
Jacobian-vector product norm as a function of *ε*. The approximated Jacobian-vector products *D*_*ξ*_*F*_b_(*ξ*)*ξ*̃ and *D*_*ξ*_*F*_tw_(*ξ*)*ξ*̃, are evaluated at a coarse bump and a coarse travelling wave *ξ* in a randomly selected direction *εξ*̃, where ||*ξ*||_2_ = 1. (a) A single realisation of the deterministic coarse-evolution maps is used in the test, showing that the norm of the Jacobian-vector product is an 𝒪(*ε*), as expected. Parameters: *p* =1, *k* = 30, *β* → ∞ (Heaviside firing rate), *h* =1, *N* = 128, *A*_1_ = 5.25, *A*_2_ = 5, *B*_1_ = 0.2, *B*_2_ = 0.3. (b) The experiment is repeated for a coarse travelling wave in the stochastic setting and for various values of *M*. Parameters as in (a), except *p* = 0.4.

We then proceed to verify directly the convergence history of the damped Newton solver. In Figure 15(a), we use a damping factor 0.5 and show the residual of the problem as a function of the number of iterations, showing that the method converges quickly to a solution. At first sight, it is surprising that the achievable tolerance of the problem does not change when the number of realisations increases. A second experiment, however, reported in Figure 15(b), shows that this behaviour is caused by the low system size: when we increase *N* from 2^7^ to 2^9^, the achievable tolerance decreases by one order of magnitude.

**Fig. 15.**
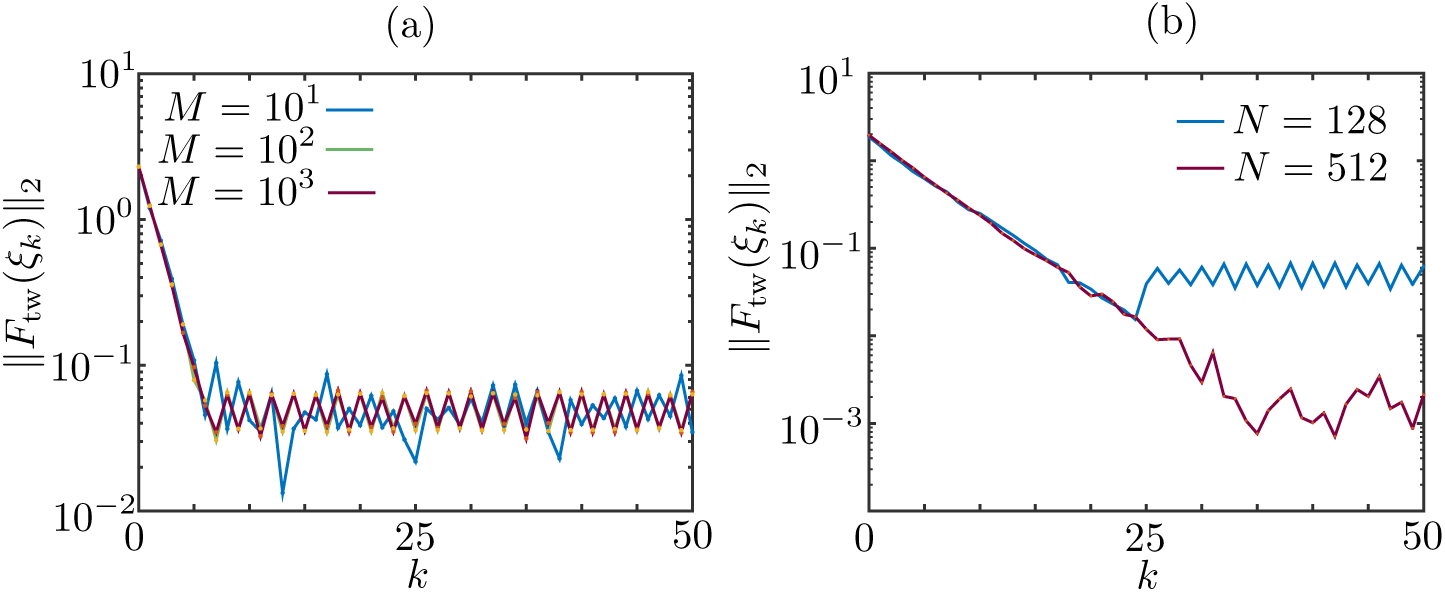
Convergence history of the damped Newton’s method applied to the coarse travelling wave problem. (a): the method converges linearly, and the achievable tolerance does not decrease when the number of realisations *M* is increased. (b): the achievable tolerance depends on the grid size, or, equivalently, on the number of neurons, *N*.

### 10.1. Numerical Bifurcation Analysis

Gong and Robinson [36], and Qi and Gong [62] found wandering spots and propagating ensembles using direct numerical simulations on the plane. Here, we perform a numerical bifurcation analysis with various control parameters for the structures found in Section 3 on a one-dimensional domain.

In Figure 16(a), we vary the primary control parameter *κ*, the gain of the convolution term, therefore, we study existence and stability of the bumps and the travelling pulse when global coupling is varied. This continuation is performed for a bump, a multiple bump and a travelling pulse in the continuum deterministic model, using Equations (5.5), (5.8)and (6.2), respectively.

**Fig. 16.**
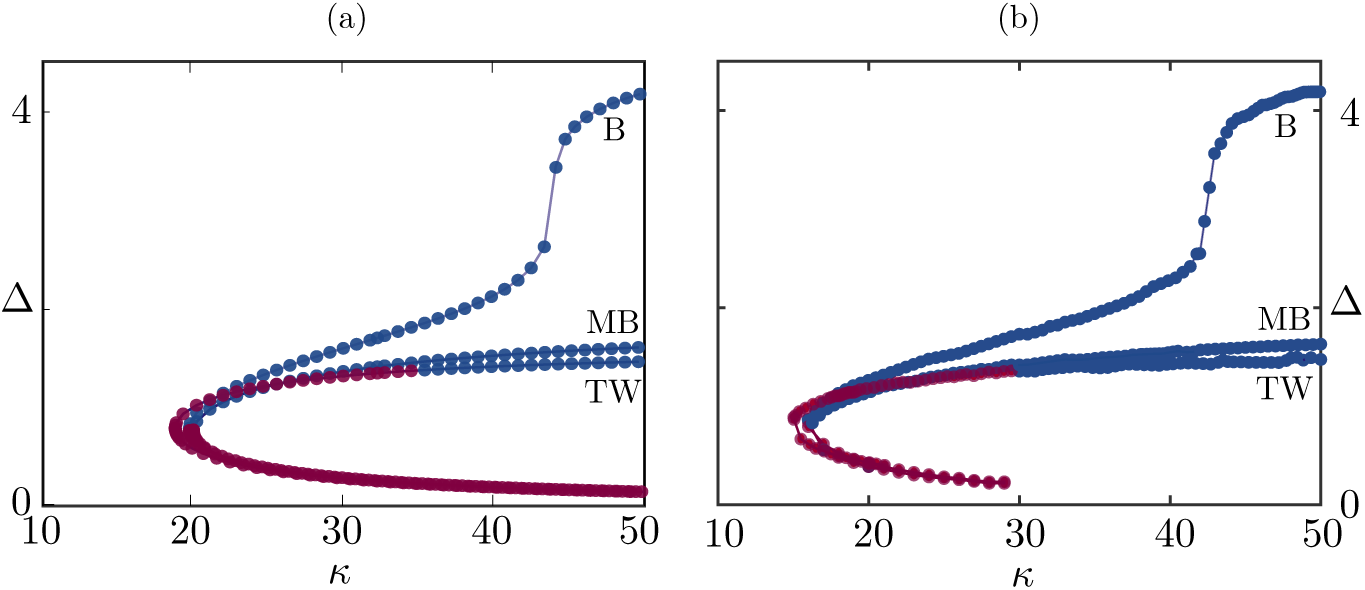
Bifurcation diagrams for bumps (B), multibumps (MB) and travelling waves (TW) using *k* as bifurcation parameter parameter, (a): Using the analytical results, we see that bump, multi-bump and travelling wave solutions coexist and are stable for sufficiently high *k* (see main text for details), (b): The solution branches found using the equation-free methods agree with the analytical results, Parameters as in Table 1 except *h* = *p* = 1.0, *β* → ∞.

For sufficiently high *k*, these states coexist and are stable in a large region of parameter space. We stress that spatially homogeneous mesoscopic states *J*(*x*) ≡ *J*_*_, with 0 = *J*_*_ or *J*_*_ > *h* are also supported by the model, but are not treated here. Interestingly, the three solution branches are disconnected, hence the bump analysed in this study does not derive from an instability of the trivial state. A narrow unstable bump Δ ≪ 1 exists for arbitrarily large *κ* (red branch); as *κ* decreases, the branch stabilises at a saddle-node bifurcation. At *κ* ≈ 42, the branch becomes steeper, the maximum of the bump changes concavity, developing a dimple. On an infinite domain, the branch displays an asymptote (not shown) as the bump widens indefinitely. On a finite domain, like the one reported in the figure, there is a maximum achievable width of the bump, due to boundary effects. The travelling wave is also initially unstable, but does not stabilise at the saddle node bifurcation. Instead, the wave becomes stable at *κ* ≈ 33, confirming the numerical simulations reported in Figure 9.

In Figure 16(b), we repeat the continuation for the same parameter values, but on a finite network, using the coarse time-steppers outlined in Sections 8.1, 8.2. The numerical procedure returns results in line with the continuum case, even at the presence of the noise induced by the finite size. The branches terminate for large *κ* and low Δ: this can be explained by noting that, if *J*(*x*) ≡ 0, then the system attains the trivial resting state *u*(*x*) ≡ 0 immediately, as no neuron can fire; on a continuum network, Δ can be arbitrarily small, hence the branch can be followed for arbitrarily large *κ*; on a discrete network, there is a minimal value of Δ that can be represented with a finite grid.

We now consider continuation of solutions in the stochastic model. In Figure 17, we vary the transition probability, *p*, from the refractory to quiescent state. In panel (a), we show analytical results, given by solving (7.5)-(7.6), whilst panel (b) shows results found using the equation-free method. We find qualitatively similar diagrams in both cases, though we note some quantitative differences, owing to the finite size of the network and the finiteness of *β*: at the presence of noise, the stationary solutions exist for a wider region of parameter space (compare the folds in Figure 17(a) and Figure 17(b)); a similar situation arises, is also valid for the travelling wave branches.

**Fig. 17.**
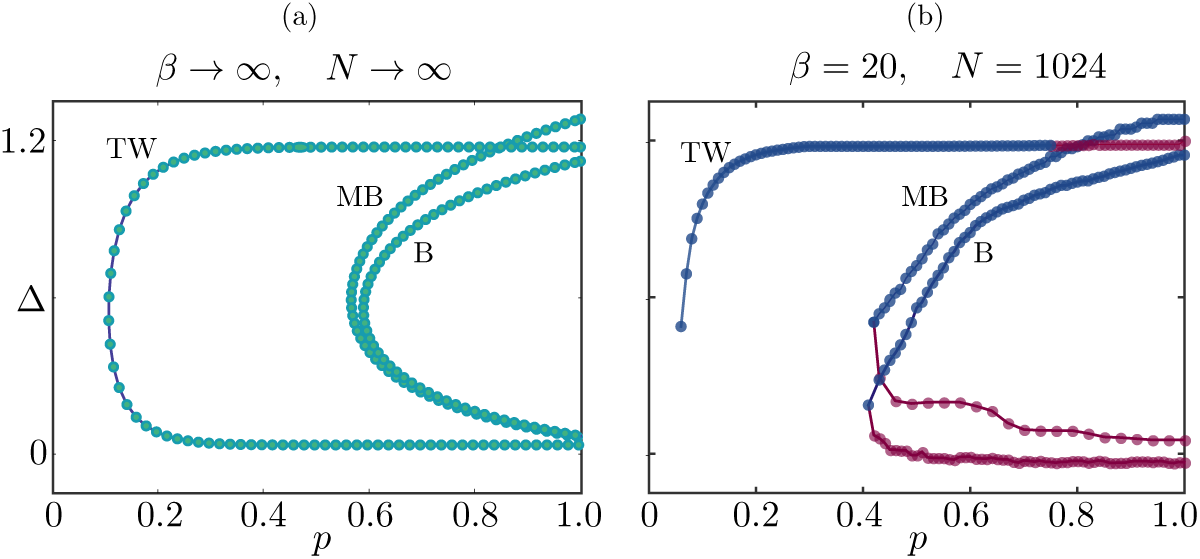
Bifurcation in the control parameter *p*, (a): Existence curves obtained analytically; we see that, below a critical value of *p*, only the travelling wave exists, (b): The solution branches found using the equation-free method agree qualitatively with the analytical results, and we can use the method to infer stability, For full details, please refer to the text, Parameters as in Table 1 except *k* = 20.0, *h* = 0.9, with *β* → ∞ for (a) and *β* = 20.0 for (b).

The analytical curves of Figure 17(a) do not contain any stability information, which are instead available in the equation-free calculations of Figure 17(b), confirming that bump and multi-bump destabilise at a saddle-node bifurcation, whereas the travelling wave becomes unstable to perturbations in the wake, if *p* is too large. The lower branch of the travelling wave is present in the analytical results, but not in the numerical ones, as this branch is not captured by our lifting strategy: when we lift a travelling wave for very low values of Δ, we have that *J* < *h* for all *x* ∈ 𝕊_*N*_ and the network attains the trivial state *u*(*x*) ≡ 0 in 1 or 2 time steps, thereby the coarse time stepper becomes ineffective, as the integration time *T* can not be reduced to 0.

Gong and co-workers [36, 62] found that refractoriness is a key component to generating propagating activity in the network. The bifurcation diagram presented here confirm this, as we recognise 3 regimes: for high *p* (low refractory time) the system supports stationary bumps, as the wave is unstable; for intermediate *p*, travelling and stationary bumps coexist and are stable, while for low *p* (high refractory time) the system selects the travelling wave.

In Figure 18, we perform the same computation now varying *β*, which governs the sensitivity of the transition from quiescence to spiking. Here, we see that the wave and both bump solutions are stable for a wide range of *β* values and furthermore, that these states are largely insensitive to variations in this parameter, implying that the Heaviside limit is a good approximation for the network in this region of parameter space.

**Fig. 18.**
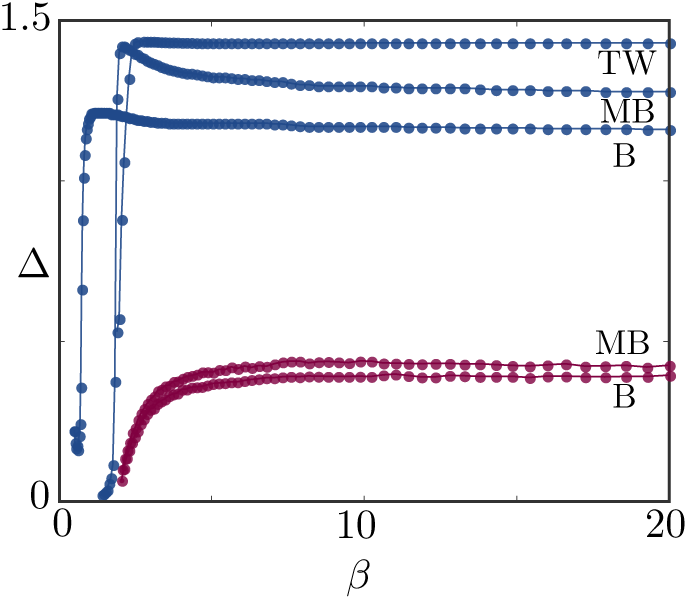
Bifurcation in the control parameter *β*, For a large range of values, we observe very little change in Δ as *β* is varied, Parameters as in Table 1 except *k* = 40.0, *h* = 0.9, *p* = 1.0, See the main text for full details.

**Fig. 19.**
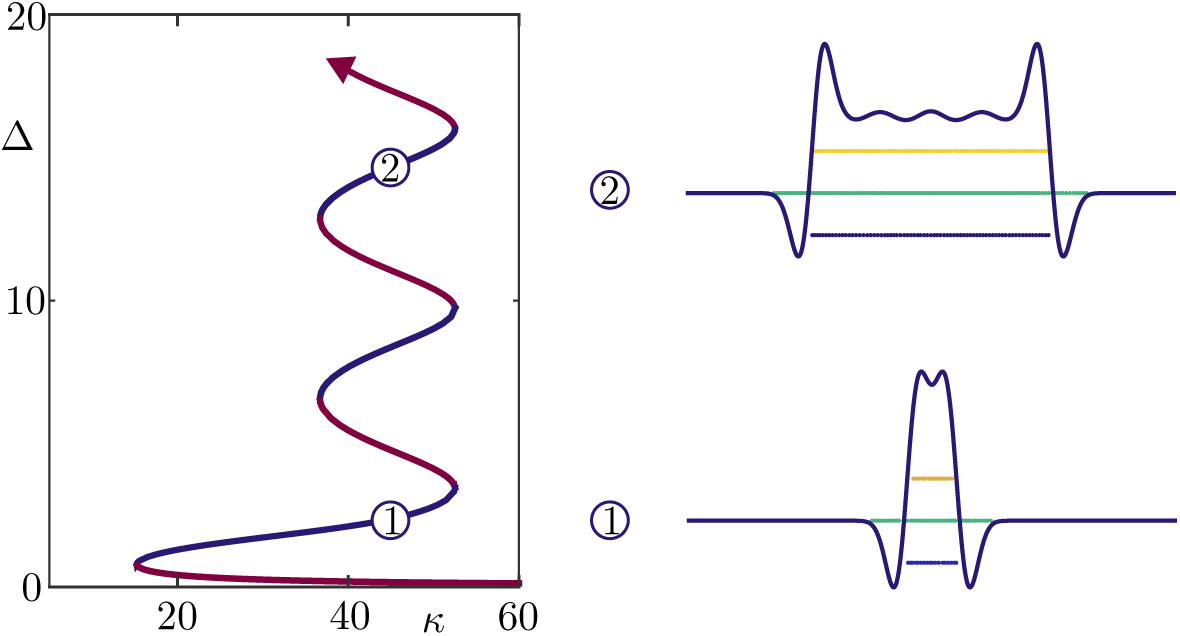
Bifurcation diagram for bumps in a heterogeneous network, To generate this figure, we replaced the coupling function with *W*̃(*x*, *y*) = *W*(*x* – *y*)(1 + *W*_0_ cos(*y*/*s*)), with *W*_0_ = 0.01, *s* = 0.5, We observe the snaking phenomenon in the approximate interval *k* ∈ [38, 52], The branches moving upwards and to the right are stable, whereas those moving to the left are unstable, The images on the right, obtained via direct simulation, depict the solution profiles on the labelled part of the branches, We note the similarity of the mesoscopic profiles within the middle of the bump, The continuation was performed for the continumm, deterministic model with parameters are *k* = 30, *h* = 0.9.

Finally, we apply the framework presented in the previous sections to study heterogeneous networks. We modulate the synaptic kernel using a harmonic function, as studied in [4] for a neural field. As in [4], the heterogeneity promotes the formation of a hierarchy of stable coexisting localised bumps, with varying width, arranged in a classical snaking bifurcation diagram. A detailed study of this bifurcation structure, while possible, is outside the scope of the present paper.

## 11. Discussion

In this article, we have used a combination of analytical and numerical techniques to study pattern formation in a Markov chain neural network model. Whilst simple in nature, the model exhibits rich dynamical behaviour, which is often observed in more realistic neural networks. In particular, spatio-temporal patterns in the form of bumps have been linked to working memory [33, 17, 34], whilst travelling waves are thought to be important for plasticity [6] and memory consolidation [57, 64]. Overall, our results reinforce the findings of [36], namely that refractoriness is key to generating propagating activity: we have shown analytically and numerically that waves are supported by a combination of high gains in the synaptic input and moderate to long refractory times. For high gains and short refractory times, the network supports localised, meandering bumps of activity.

The analysis presented here highlights the multiscale nature of the model by showing how evolution on a microscopic level gives rise to emergent behaviour at meso‐ and macroscopic levels. In particular, we established a link between descriptions of the model at multiple spatial scales: the identified coarse spatiotemporal patterns have typified and recognisable motifs at the microscopic level, which we exploit to compute macroscopic patterns and their stability.

Travelling waves and bumps have almost identical meso‐ and macroscopic profiles: if microscopic data were removed from Figure 2(a) and Figure 4(a), the profiles and activity sets of these two patterns would be indistinguishable. We have shown that a disambiguation is however possible if the meso‐ and macroscopic descriptions take into account microscopic traits of the patterns: in the deterministic limit of the system, where mathematical analysis is possible, the microscopic structure is used in the partition sets of Propositions 5.1 and 6.1; in the stochastic setting with Heaviside firing rates and infinite number of neurons, the microscopic structure is reflected in the approximate probability mass functions appearing in Section 7; in the full stochastic finite-size setting, where an analytical description is unavailable, the microscopic structure is hardwired in the lifting operators of the coarse time-steppers (Section 8).

An essential ingredient in our analysis is the dependence of the Markov chain transition probability matrix upon the global activity of the network, via the firing rate function *f*. Since this hypothesis is used to construct rate models as Markov processes [10], our lifting strategy could be used in equation-free schemes for more general large-dimensional neural networks. An apparent limitation of the procedure presented here is its inability to lift strongly unstable patterns with low activity, as pointed out in Section 10. This limitation, however, seems to be specific to the model studied here: when Δ → 0, bumps destabilise with transients that are too short to be captured by the coarse time-stepper.

A possible remedy would be to represent the pattern via a low-dimensional, spatially-extended, spectral discretisation of the mesoscopic profile (see [48]), which would allow us to represent the synaptic activity below the threshold *h*. This would lead to a larger-dimensional coarse system, in which noise would pollute the Jacobian-vector evaluation and the convergence of the Newton method. Variance-reduction techniques [65] have been recently proposed for equation-free methods in the context of agent-based models [3], and we aim to adapt them to large neural networks in subsequent publications.

## Acknowledgments

We are grateful to Joel Feinstein, Gabriel Lord and Wilhelm Stannat for helpful discussions and comments on a preliminary draft of the paper. Kyle Wedgwood was generously supported by the Wellcome Trust Institutional Strategic Support Award (WT105618MA)

## REFERENCES

[1] Shun-ichi Amari. Homogeneous nets of neuron-like elements. Biological Cybernetics, 17(4):211–220, 1975.

[2] Shun-ichi Amari. Dynamics of pattern formation in lateral-inhibition type neural fields. Biological Cybernetics, 27(2):77–87, June 1977.

[3] Daniele Avitabile, Rebecca Hoyle, and Giovanni Samaey. Noise reduction in coarse bifurcation analysis of stochastic agent-based models: an example of consumer lock-in. SIAM Journal on Applied Dynamical Systems, 13(4):1583–1619, 2014.

[4] Daniele Avitabile and Helmut Schmidt. Snakes and ladders in an inhomogeneous neural field model. Physica D: Nonlinear Phenomena, 294:24–36, 2015.

[5] Javier Baladron, Diego Fasoli, Olivier Faugeras, and Jonathan Touboul. Mean-field description and propagation of chaos in networks of Hodgkin-Huxley and FitzHugh-Nagumo neurons. The Journal of Mathematical Neuroscience, 2(10), 2012.

[6] James E. M. Bennett. Refinement and pattern formation in neural circuits by the interaction of traveling waves with spike-timing dependent plasticity. PLoS Computational Biology, 11(8):e1004422, 2015.

[7] Chris A. Brackley and Matthew S. Turner. Random fluctuations of the firing rate function in a continuum neural field model. Physical Review E, 75(4):041913, 2007.

[8] Valentino Braitenberg and Almut Schiiz. Cortex: Statistics and Geometry of Neuronal Connectivity. Springer Berlin Heidelberg, Berlin, Heidelberg, 1998.

[9] Paul C. Bressloff. Stochastic neural field theory and the system-size expansion. SIAM Journal on Applied Mathematics, 70(5):1488–1521, 2009.

[10] Paul C. Bressloff. Stochastic Neural Field Theory and the System-Size Expansion. SIAM Journal on Applied Mathematics, 70(5):1488–1521, January 2010.

[11] Paul C. Bressloff. Spatiotemporal dynamics of continuum neural fields. Journal of Physics A: Mathematical and Theoretical, 45(3):033001, 2012.

[12] Paul C. Bressloff. Waves in Neural Media. Lecture Notes on Mathematical Modelling in the Life Sciences. Springer New York, New York, NY, 2014.

[13] Paul C. Bressloff, Jack D. Cowan, Martin Golubitsky, and Peter J. Thomas. Scalar and pseu doscalar bifurcations motivated by pattern formation on the visual cortex. Nonlinearity, 14(4):739–775, July 2001.

[14] Paul C. Bressloff and Zachary P. Kilpatrick. Two-dimensional bumps in piecewise smooth neural fields with synaptic depression. SIAM Journal on Applied Mathematics, 71(2):379–408, 2011.

[15] Paul C. Bressloff and Matthew A. Webber. Front propagation in stochastic neural fields. SIAM Journal on Applied Dynamical Systems, 11(2):708–740, 2012.

[16] David Cai, Louis Tao, Michael Shelley, and David W. McLaughlin. An effective kinetic representation of fluctuation-driven neuronal networks with application to simple and complex cells in visual cortex. Proceedings of the National Academy of Sciences, 101(20):7757–7762, May 2004.

[17] C.L. Colby, J.R. Duhamel, and M.E. Goldberg. Oculocentric spatial representation in parietal cortex. Cereb. Cortex, 5:470–481, 1995.

[18] Stephen Coombes and Markus R. Owen. Evans functions for integral neural field equations with heaviside firing rate function. SIAM Journal on Applied Dynamical Systems, 3(4):574–600, 2004.

[19] Stephen Coombes, Helmut Schmidt, and Ingo Bojak. Interface dynamics in planar neural field models. Journal of Mathematical Neuroscience, 2(9), 2012.

[20] Stephne Coombes, Helmut Schmidt, and Daniele Avitabile. Neural Field Theory, chapter Spots: Breathing, drifting and scattering in a neural field model. Springer, 2013.

[21] Weinan E, Bjorn Engquist, Xiantao Li, Weiqing Ren, and Eric Vanden-Eijden. Heterogeneous multiscale method: a review. Communications in Computational Physics, 2:367–450, 2007.

[22] Gaute T Einevoll, Christoph Kayser, Nikos K. Logothetis, and Stefano Panzeri. Modelling and analysis of local field potentials for studying the function of cortical circuits. Nature Reviews Neuroscience, 14(11):770–785, November 2013.

[23] G B Ermentrout and J D Cowan. A mathematical theory of visual hallucination patterns. Biological Cybernetics, 34(3):137–150, 1979.

[24] G. Bard Ermentrout. Neural networks as spatio-temporal pattern-forming systems. Reports on Progress in Physics, 61(4):353–430, April 1998.

[25] G. Bard Ermentrout and J. Bryce McLeod. Existence and uniqueness of travelling waves for a neural network. Proceedings of the Royal Society of Edinburgh: Section A Mathematics, 123:461–478, 1993.

[26] G. Bard Ermentrout and David H. Terman. Mathematical Foundations of Neuroscience. Springer New York, New York, NY, 2010.

[27] Adrienne Fairhall and Haim Sompolinsky. Editorial overview: theoretical and computational neuroscience. Current Opinion in Neurobiology, 25:v–viii, April 2014.

[28] Olivier Faugeras, Jonathan Touboul, and Bruno Cessac. A constructive mean-field analysis of multi-population neural networks with random synaptic weights and stochastic inputs. Frontiers in Computational Neuroscience, 3:1–28, 2009.

[29] Gregory Faye, James Rankin, and David J. B. Lloyd. Localized radial bumps of a neural field equation on the Euclidean plane and the Poincare disk. Nonlinearity, 26:437–478, 2013.

[30] Stefanos E. Folias and Paul C. Bressloff. Breathing pulses in an excitatory neural network. SIAM Journal on Applied Dynamical Systems, 3(3):378–407, 2004.

[31] Stefanos E. Folias and Paul C. Bressloff. Breathers in two-dimensional neural media. Physical Review Letters, 95(20):208107, 2005.

[32] Stefanos E. Folias and G. Bard Ermentrout. Bifurcations of Stationary Solutions in an Interacting Pair of E-I Neural Fields. SIAM Journal on Applied Dynamical Systems, 11(3):895–938, August 2012.

[33] S. Funahashi, C.J. Bruce, and P.S. Goldman-Rakic. Mnemonic coding of visual space in the monkey’s dorsolateral prefrontal cortex. Journal of Neurophysiology, 61:331–349, 1989.

[34] P.S. Goldman-Rakic. Cellular basis of working memory. Neuron, 14:477–485, 1995.

[35] David Golomb and G. Bard Ermentrout. Continuous and lurching traveling pulses in neuronal networks with delay and spatially decaying connectivity. Proceedings of the National Academy of Sciences, 96(23):13480–13485, November 1999.

[36] Pulin Gong and Peter A. Robinson. Dynamic pattern formation and collisions in networks of excitable elements. Physical Review E, 85(5):055101(R), 2012.

[37] E. Haskell, D. Q. Nykamp, and D. Tranchina. A population density method for large-scale modeling of neuronal networks with realistic synaptic kinetics. Neurocomputing, 38-40:627–632, June 2001.

[38] Axel Hutt, Andre Longtin, and Lutz Schimansky-Geier. Additive noise-induced turing transitions in spatial systems with application to neural fields and the swift—hohenberg equation. Physica D: Nonlinear Phenomena, 237(6):755–773, 2008.

[39] Eugene M. Izhikevich. Dynamical systems in neuroscience. MIT Press, 2007.

[40] Eugene M. Izhikevich and Edelman Gerald M. Large-scale model of mammalian thalamocortical systems. Proceedings of the National Academy of Sciences, 105(9):3593–3598, March 2008.

[41] Viktor K. Jirsa and Hermann Haken. A derivation of a macroscopic field theory of the brain from the quasi-microscopic neural dynamics. Physica D: Nonlinear Phenomena, 99(4):503–526, January 1997.

[42] Ioannis G. Kevrekidis, C. William Gear, James M. Hyman, Panagiotis G. Kevrekidis, Olof Runborg, and Constantinos Theodoropoulos. Equation-free, coarse-grained multiscale computation: enabling microscopic simulators to perform system-level tasks. Communications in Mathematical Sciences, 1(4):715–762, 2003.

[43] Ioannis G Kevrekidis and Giovanni Samaey. Equation-Free Multiscale Computation: Algorithms and Applications. Annual Review of Physical Chemistry, 60(1):321–344, May 2009.

[44] Zachary P. Kilpatrick and Paul C. Bressloff. Stability of bumps in piecewise smooth neural fields with nonlinear adaptation. Physica D: Nonlinear Phenomena, 239(12):1048–1060, 2010.

[45] Zachary P. Kilpatrick and G. Bard Ermentrout. Wandering bumps in stochastic neural fields. SIAM Journal on Applied Dynamical Systems, 12(1):61–94, 2013.

[46] Christian Kuehn and Martin Riedler. Large deviations for nonlocal stochastic neural fields. Journal Mathematical Neuroscience, 4(1):1–33, 2014.

[47] Carlo R. Laing. Spiral waves in nonlocal equations. SIAM Journal on Applied Dynamical Systems, 4(3):588–606, 2005.

[48] Carlo R. Laing. On the application of “equation-free modelling” to neural systems. Journal of Computational Neuroscience, 20(1):5–23, February 2006.

[49] Carlo R Laing, Thomas Frewen, and Ioannis G Kevrekidis. Reduced models for binocular rivalry. Journal of Computational Neuroscience, 28(3):459–476, 2010.

[50] Carlo R. Laing, Thomas A. Frewen, and Ioannis G. Kevrekidis. Coarse-grained dynamics of an activity bump in a neural field model. Nonlinearity, 20(9):2127–2146, September 2007.

[51] Carlo R Laing and Ioannis G Kevrekidis. Equation-free analysis of spike-timing-dependent plasticity. Biological Cybernetics, 109(6):701–714, 2015.

[52] Carlo R. Laing and William C. Troy. PDE methods for nonlocal models. SIAM Journal on Applied Dynamical Systems, 2(3):487–516, 2003.

[53] Carlo R. Laing and William C. Troy. PDE Methods for Nonlocal Models. SIAM Journal on Applied Dynamical Systems, 2(3):487–516, 2003.

[54] Carlo R. Laing, William C. Troy, Boris Gutkin, and G. Bard Ermentrout. Multiple bumps in a neuronal model of working memory. SIAM Journal on Applied Mathematics, 63(1):62–97, 2002.

[55] Cheng Ly and Daniel Tranchina. Critical Analysis of Dimension Reduction by a Moment Closure Method in a Population Density Approach to Neural Network Modeling. Neural Computation, 19(8):2032–2092, June 2007.

[56] Henry Markram, Eilif Muller, Srikanth Ramaswamy, Michael W. Reimann, Marwan Abdellah, Carlos Aguado Sanchez, Anastasia Ailamaki, Lidia Alonso-Nanclares, Nicolas Antille, Selim Arsever, Guy Antoine Atenekeng Kahou, Thomas K. Berger, Ahmet Bilgili, Nenad Buncic, Athanassia Chalimourda, Giuseppe Chindemi, Jean Denis Courcol, Fabien Delalondre, Vincent Delattre, Shaul Druckmann, Raphael Dumusc, James Dynes, Stefan Eilemann, Eyal Gal, Michael Emiel Gevaert, Jean-Pierre Ghobril, Albert Gidon, Joe W. Graham, Anirudh Gupta, Valentin Haenel, Etay Hay, Thomas Heinis, Juan B. Hernando, Michael Hines, Lida Kanari, Daniel Keller, John Kenyon, Georges Khazen, Yihwa Kim, James G. King, Zoltan Kisvarday, Pramod Kumbhar, Sebastien Lasserre, Jean Vincent Le Be, Bruno R. C. Magalhaes, Angel Merchan-Perez, Julie Meystre, Benjamin Roy Morrice, Jeffrey Muller, Alberto Munoz-Cespedes, Shruti Muralidhar, Keerthan Muthurasa, Daniel Nachbaur, Taylor H. Newton, Max Nolte, Aleksandr Ovcharenko, Juan Palacios, Luis Pastor, Rodrigo Perin, Rajnish Ranjan, Imad Riachi, Jose-Rodrigo Rodriguez, Juan Luis Riquelme, Christian Rossert, Konstantinos Sfyrakis, Ying Shi, Julian C. Shillcock, Gilad Silberberg, Ricardo Silva, Farhan Tauheed, Martin Telefont, Maria ToledoRodriguez, Thomas Trankler, Werner Van Geit, Jafet Villafranca Diaz, Richard Walker, Yun Wang, Stefano M. Zaninetta, Javier Defelipe, Sean L. Hill, Idan Segev, and Felix Schurmann. Reconstruction and simulation of neocortical microcircuitry. Cell, 163:456492, 2015.

[57] Marcelo Massimini, Reto Huber, Fabio Ferrarelli, Sean Hill, and Giulio Tononi. The sleep slow oscillation as a traveling wave. Journal of Neuroscience, 4(24):6862–6870, 2004.

[58] Paul L. Nunez and Ramesh Srinivasan. Electric fields of the brain: the neurophysics of EEG. Oxford University Press, 2nd edition, 2006.

[59] Ahmet Omurtag, Bruce W. Knight, and Lawrence Sirovich. On the Simulation of Large Populations of Neurons. Journal of Computational Neuroscience, 8(1):51–63, 2000.

[60] Remus Osan and Bard Ermentrout. Two dimensional synaptically generated traveling waves in a theta-neuron neural network. Neurocomputing, 38-40:789–795, 2001.

[61] Markus R. Owen, Carlo R. Laing, and Stephen Coombes. Bumps and rings in a two-dimensional neural field: splitting and rotational instabilities. New Journal of Physics, 9(10):378–401, 2007.

[62] Yang Qi and Pulin Gong. Dynamic patterns in a two-dimensional neural field with refractoriness. Physical Review E, 92(2):022702, 2015.

[63] James Rankin, Daniele Avitabile, Javier Baladron, Gregory Faye, and David J B Lloyd. Continuation of Localized Coherent Structures in Nonlocal Neural Field Equations. SIAM Journal on Scientific Computing, 36(1):B70–B93, January 2014.

[64] Bjorn Rasch and Jan Born. About sleep’s role in memory. Physiological Reviews, 93(2):681–766, 2013.

[65] Mathias Rousset and Giovanni Samaey. Simulating individual-based models of bacterial chemotaxis with asymptotic variance reduction. Mathematical Models and Methods in Applied Sciences, 23(12):2155–2191, August 2013.

[66] Konstantinos G. Spiliotis and Constantinos I. Siettos. A timestepper-based approach for the coarse-grained analysis of microscopic neuronal simulators on networks: Bifurcation and rare-events micro-to macro-computations. Neurocomputing, 74(17):3576–3589, 2011.

[67] Konstantinos G. Spiliotis and Constantinos I. Siettos. Multiscale computations on neural networks: from the individual neuron interactions to the macroscopic-level analysis. International Journal of Bifurcation and Chaos, 20(01):121–134, May 2012.

[68] Laurette S Tuckerman and Dwight Barkley. Bifurcation Analysis for Timesteppers. In Numerical Methods for Bifurcations of Dynamical Equilibria, pages 453–466. SIAM, New York, NY, January 2000.

[69] Martijn P. van den Heuvel and Hilleke E. Hulshoff Pol. Exploring the brain network: A review on resting-state fMRI functional connectivity. European Neuropsychopharmacology, 20(8):519–534, August 2010.

[70] Weinan W and Bjorn Engquist. The heterogeneous multiscale methods. Communications in Mathematical Sciences, 1(1):87–132, 2003.

[71] Thomas M. Wasylenko, Jaime E. Cisternas, Carlo R. Laing, and Ioannis G. Kevrekidis. Bifurcations of lurching waves in a thalamic neuronal network. Biological Cybernetics, 103(6):447–462, 2010.

[72] Herrad Werner and Tim Richter. Circular stationary solutions in two-dimensional neural fields. Biological Cybernetics, 85(3):211–217, September 2001.

[73] Hugh R. Wilson and Jack D. Cowan. Excitatory and inhibitory interactions in localized populations of model neurons. Biophysics Journal, 12:1–24, 1972.

[74] Hugh R. Wilson and Jack D. Cowan. A mathematical theory of the functional dynamics of cortical and thalamic nervous tissue. Biological Cybernetics, 13(2):55–80, September 1973.

[75] Rafael Yuste. From the neuron doctrine to neural networks. Nature Reviews Neuroscience, 16(8):487–497, 2015.

